# Comprehensive longitudinal profiling of SARS-CoV-2-specific CD8^+^ T-cells reveal strong functional impairment and recognition bias as markers for disease severity

**DOI:** 10.1101/2025.03.25.644515

**Authors:** Susana Patricia Amaya Hernandez, Kamilla Kjærgaard Munk, Konstantin Danilov, Mohammad Kadivar, Ditte Stampe Hersby, Tripti Tamhane, Simone Majken Stegenborg-Grathwohl, Anne Ortved Gang, Sine Reker Hadrup, Sunil Kumar Saini

## Abstract

CD8^+^ T-cells are essential for controlling and resolving SARS-CoV-2 infection, yet their antigen-specific resolution in relation to disease severity, functional dynamics during acute infection, and long-term memory formation remain incompletely understood. Using comprehensive longitudinal profiling of 553 SARS-CoV-2 immunogenic antigens across globally prevalent HLAs, we identified antigen-specific CD8^+^ T-cell responses that were either critical for early viral clearance or associated with severe disease outcomes.

During acute infection, patients with severe COVID-19 exhibited a broader and more robust CD8^+^ T-cell response than those with mild disease. Notably, we identified HLA-A1-restricted immunodominant antigen-specific T-cells strongly associated with severe disease. These T-cells were present at extremely high frequencies but showed significantly reduced expression of cytotoxic molecules at both the transcriptomic (*PRF1, GZMB, GZMH, GNLY*) and protein levels (IFN-γ, TNF-α, IL-2), as revealed by multidimensional single-cell and cytokine profiling. In contrast, patients with mild disease had T-cells that recognized a more restricted set of antigens, showed only partial overlap with those in severe cases, and showed enhanced cytotoxicity, along with enrichment in gene sets associated with cytotoxic function, hypoxia, and glycolysis. Furthermore, the long-term memory CD8^+^ T-cells were maintained for a limited subset of immunodominant antigens, with their persistence correlating with their initial frequency during infection. Importantly, SARS-CoV-2 vaccination following infection expanded the long-term T-cell repertoire by enhancing pre-existing responses and generating de novo responses, regardless of prior disease severity.

These findings resolve the antigen-specific kinetics and durability of CD8^+^ T-cells in SARS-CoV-2 infection and provide key insights into their functional landscape. This knowledge could inform future vaccine strategies and therapeutic interventions to enhance protective immunity against emerging viral threats.

## Introduction

Since the emergence of severe acute respiratory syndrome coronavirus 2 (SARS-CoV-2), more than 7 million deaths and over 774 million confirmed cases have been reported globally as a result of this novel virus (1). COVID-19, the disease caused by SARS-CoV-2, presents a broad spectrum of clinical manifestations, ranging from asymptomatic or mild flu-like symptoms, with patients recovering without hospitalization, to severe pneumonia, acute respiratory distress syndrome (ARDS), and even multi-organ failure, often requiring intensive care and mechanical ventilation (2–4). The clinical outcomes of COVID-19 are influenced by a combination of risk factors, including age, sex, underlying comorbidities (such as hypertension, diabetes, chronic pulmonary, kidney, or liver disease, immunodeficiencies, cancer, cardiovascular disease, and obesity), and genetic predispositions, which can affect both the viral pathogenesis and the host immune response (2,5). The variable nature of these clinical outcomes emphasizes the critical need for a deeper understanding of the immune responses to SARS-CoV-2 infection and the associated disease outcome.

The role of CD8^+^ T-cells has been well documented in mild and asymptomatic SARS-CoV-2 infection and protection from severe COVID-19. This is particularly significant considering the observed decline in antibody responses over time (6–9) and lesser protection against evolving SARS-CoV-2 variants (10,11). Conversely, dysfunction of CD8^+^ T-cells has also been reported in severe COVID-19, which includes upregulation of exhaustion markers and impaired cytokine production (12,13). Despite the potential dysfunction, we and others have shown robust T-cell activation and a higher magnitude of CD8^+^ T-cell response in severe COVID-19 patients (9,14,15). Such differential T-cell response and disease association requires a large-scale analysis resolving antigen-specific CD8^+^ T-cells at the early onset of the infection and the influence of such T-cells in establishing long-term memory. However, due to the novel nature of SARS-CoV-2, the antigen-specific immune response and its impact on both short-and long-term immune response remain inadequately understood.

We have previously shown that in the acute phase of SARS-CoV-2 infection, CD8^+^ T-cell response is generated against a substantially large fraction of SARS-CoV-2 antigens, and a higher frequency of antigen-specific T-cells was observed in hospitalized patients compared to patients with mild disease (14). In our analysis of >3000 peptides, we reported 424 immunogenic peptides identified in COVID-19 patients and samples from healthy donors. Similarly, other large-scale analyses of COVID-19 patients have identified a broad range of immunogenic epitopes, constituting more than 1000 CD8^+^ T-cell epitopes in the Immune Epitope Database (IEDB) data repository, covering also variants of SARS-CoV2 (16). The kinetics of CD8^+^ T-cells in longitudinal analysis ranging from a few weeks to several months has been reported by a few studies (17–20). However, most of the longitudinal analysis followed a few antigen-specific T-cells or performed analysis based on peptide pools, thus providing limited epitope-specific resolution.

In this study, we utilized DNA-barcoded peptide-MHC (pMHC) multimers in combination with a T-cell phenotype panel for a comprehensive longitudinal profiling of 553 SARS-CoV-2-derived CD8^+^ T-cell epitopes in patients with mild and severe COVID-19, up to approximately 220 days post-infection. Our approach allowed the precise identification and tracking of SARS-CoV-2–specific CD8^+^ T-cells at the antigen-specific level, enabling us to explore the dynamics of the immune response across different disease severities and to examine the functional and phenotypic differences in CD8^+^ T-cells between these patient groups. To further analyze the immune response, we conducted single-cell profiling of CD8^+^ T-cells targeting SARS-CoV-2 epitopes of interest. By integrating single-cell RNA sequencing, T-cell receptor sequencing, and CITE-seq antibodies, we detailed the phenotypes and functions of specific immune cell subsets, investigating their correlation with COVID-19 severity. Additionally, our study extended to analyze the T-cell response in SARS-CoV-2 infected patients who were later vaccinated with a COVID-19 mRNA vaccine. This part of the research aimed to evaluate the effect of vaccination on the T-cell repertoire of previously infected individuals, providing insights into how prior infection influences immune memory and vaccine efficacy.

## Results

### SARS-CoV-2 immunogenic antigens for longitudinal profiling of CD8**^+^** T-cells

Previously, in a study of genome-wide mapping of SARS-CoV-2 immunogenic CD8^+^ T-cell antigens, we have shown a stronger and antigen-specific immunodominance of T-cell activation in the acute phase of COVID-19 disease in severe patients compared to patients with mild COVID-19 disease (14). Now to investigate the impact and association of antigen-specific CD8^+^ T-cells in severe and mild COVID-19 disease on long-term T-cell memory and functionality, we analyzed a cohort of 73 patients infected during the first wave of the COVID-19 pandemic, over a period of approximately 220 days. This cohort comprised 36 patients classified as having severe disease and 37 with mild disease based on their requirement for hospitalization or not (see Materials and Methods for stratification details). Following the initial PCR-confirmed SARS-CoV-2 infection (April-May 2020) (Supplementary Table 1), blood samples were collected at four different time points (TPs): first collection as close as possible to the positive COVID-19 test (TP1, 9 ± 7 days post-diagnosis), and subsequently at 27 ± 10 days (TP2), 58 ± 12 days (TP3), and 218 ± 30 days (TP4) until January 2021. Within this group, 17 individuals (severe, n = 10; mild, n = 7) were vaccinated with one or two doses of a COVID-19 mRNA vaccine between TP3 and TP4 (For an overview of the study design see Fig. 1A, Supplementary Fig. 1 and Supplementary Table 2).

**Fig. 1.**
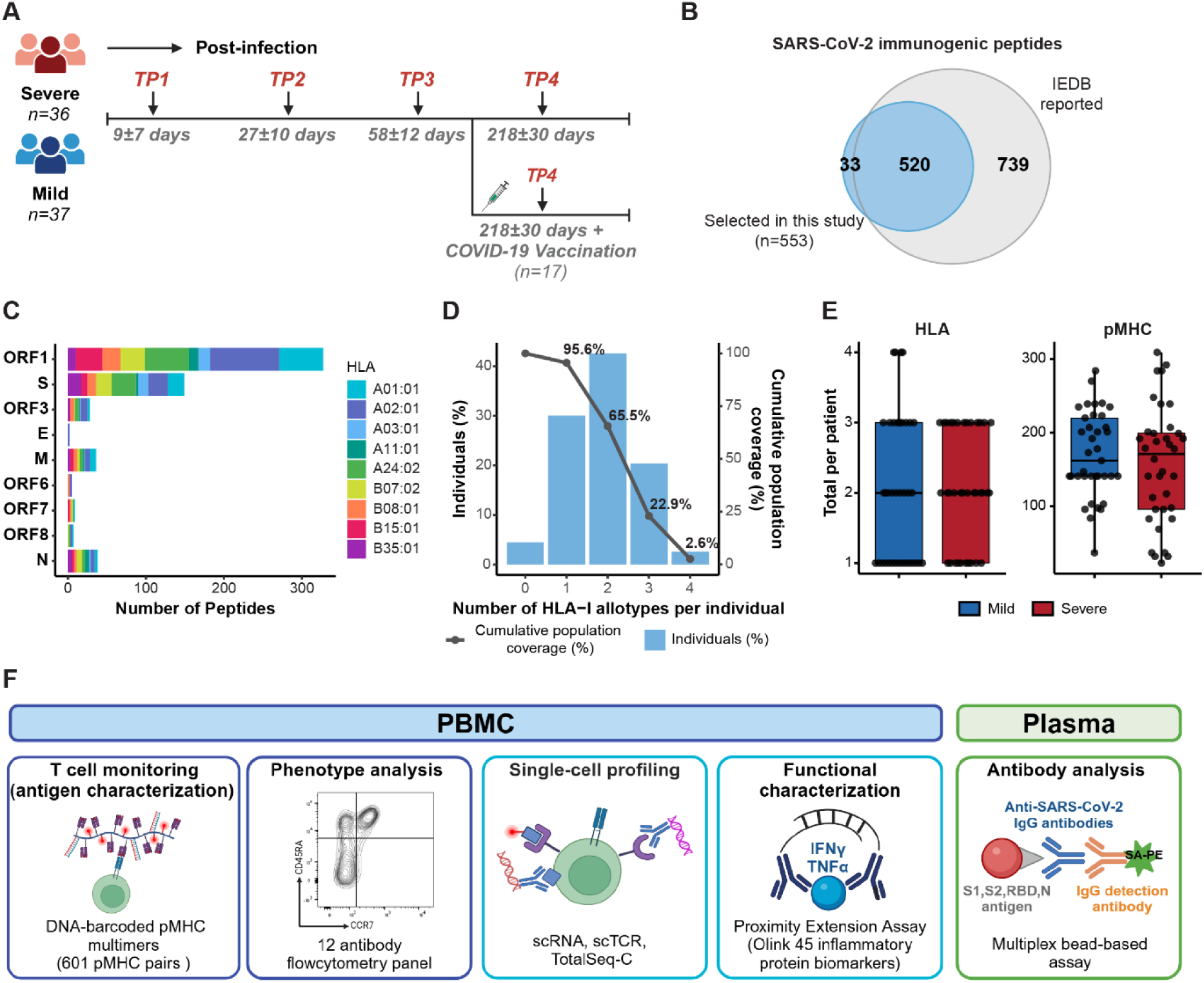
Study design and selection of SARS-CoV-2 immunogenic peptides for longitudinal analysis. (**A**) Sample collection timeline for patients with COVID-19 (Severe, n = 36 individuals; Mild, n = 37 individuals; TP: time point). Vaccinated patients received 1 or 2 doses of a COVID-19 vaccine (n = 17). (**B**) Venn diagram shows the number of SARS-CoV-2 derived epitopes included in this study (n = 553) out of the total SARS-CoV-2 epitopes reported in IEDB. (**C**) Distribution of the selected SARS-CoV-2 epitopes among the structural proteins (S, M, E, N) and non-structural proteins (ORF1, ORF3, ORF6, ORF7, and ORF8) according to their HLA-restriction. (**D**) Population coverage achieved with the selection of nine common HLA class I allotypes for SARS-CoV-2 T-cell epitope screening and analysis. The percentage of individuals within the world population carrying up to four HLA-I allotypes (*x-*axis) are indicated as blue bars on the left *y-*axis. The cumulative percentage of population coverage is depicted as grey dots on the right *y-*axis. (**E**) Comparison of the total number of HLA (left) and pMHC (right) included for analysis per patient between the severe and mild groups. Each dot represents one patient. Mann-Whitney test, mild vs severe HLA (p = 0.718), mild vs severe pMHC (p = 0.456). (**F**) Schematic diagram illustrating the different assays used in this study.

For comprehensive coverage of immunogenic CD8^+^ T-cell antigens, allowing us to differentiate T-cell repertoire and functional consequences in mild and severe COVID-19 disease, we selected a total of 553 unique peptides based on immunogenicity reported by us and others (Fig. 1B and Supplementary Table 3). Of these, 311 epitopes were identified in our previous study (Saini *et al.*, 2021 (14)). An additional 33 peptides were included based on genome-wide T-cell mapping of 19 COVID-19 patients sampled at TP1, using DNA-barcoded pMHC multimers (Supplementary Fig. 3A). In this analysis, 2,198 peptides were predicted to bind at least one of seven selected HLA alleles (HLA-A*01:01,-A*02:01,-A*03:01,-A*24:02,-B*07:02,-B*08:01, and –B*15:01) using NetMHCpan-4.1 with a rank threshold of <1% (21). These predicted peptides were subsequently screened for CD8^+^ T-cell reactivity (Supplementary Table 4). The remaining peptides were selected from IEDB based on immunogenic epitopes reported by other studies and restricted to the HLA alleles included in this study with a rank threshold of <1% predicted using NetMHCpan-4.1 (16,21).

Altogether, our peptide library for longitudinal analysis comprised a total of 553 unique epitopes representing nine prevalent HLA alleles and spanning across nine SARS-CoV-2 proteins (Fig. 1C and Supplementary Tables 3 and 5), ensuring at least one HLA allotype was represented in 95.6% of the global population (Fig. 1D). Based on the HLA types of our study cohort, we achieved an average HLA coverage of 2.08 alleles per individual in severe patients and 2.05 alleles in the mild patients (Fig. 1E and Supplementary Table 6).

The 553 peptides were used to construct 601 peptide-HLA pairs for experimental assessment (Supplementary Table 3). DNA-barcoded multimers were prepared by loading HLA-specific peptides on MHC molecules, followed by multimerization on a PE (phycoerythrin)– or APC (allophycocyanin)– labeled dextran backbone tagged with unique DNA barcodes (22).

Peripheral blood mononuclear cells (PBMCs) from each patient were incubated with HLA-matching pMHC multimers (Supplementary Table 6) and stained with a phenotype antibody panel (Supplementary Table 7) to identify multimer-reactive CD8^+^ T-cells (Fig. 1F, Supplementary Fig. 2). We evaluated on average 159 ± 79.8 and 172 ± 59.5 pMHC multimers per individual for severe and mild patients respectively (Fig. 1E). Additionally, for comparisons between SARS-CoV-2 and other well-characterized viral antigens, 38 peptides derived from cytomegalovirus (CMV), Epstein-Barr virus (EBV), and influenza (FLU), collectively referred to as CEF, were included in the multimer panel (Supplementary Table 8).

### Enhanced frequency and long-term persistence of antigen-specific T-cells in severe COVID-19

Longitudinal profiling of antigen-specific CD8^+^ T-cells identified the distribution of T-cell responses in mild and severe disease conditions in the acute phase of infection and the breadth and frequency of T-cell repertoire that established the long-term T-cell memory (Fig. 2). In the acute phase of infection at TP1 76.5% (26/34) of the severe and 75% (27/36) of the mild patients mounted CD8^+^ T-cell responses recognizing at least one of the SARS-CoV-2 peptides included in the study and remained long-term in 65.2% (15/23) of mild and 62.9% (22/35) of the severe patients at TP4 (Supplementary Table 9). In total, we identified T-cell responses to 215 pMHC complexes, across all time points and patients, corresponding to 210 unique SARS-CoV-2 T-cell epitopes across the 9 analyzed HLAs and 9 analyzed proteins (Fig. 2, Supplementary Fig. 3B, and Supplementary Table 10). Interestingly, in the early phase (TP1 and TP2), 101 pMHC complexes were detected exclusively in the severe patients, 60 exclusively in the mild patients, and only 38 were shared between the two groups, showing a broader repertoire of responses in the severe patients (Fig. 2, Fig. 3A). Furthermore, the frequency of antigen-specific T-cells and the number of unique T-cell responses were much higher in severe patients (Fig. 3B, 3C). In the acute phase of infection, severe patients had a mean overall frequency of total antigen-specific CD8^+^ T-cells (calculated as the sum of estimated frequencies for all SARS-CoV-2–specific T-cells) of 4.9 ± 9.1%. In contrast, in the mild patients, it was 1.4 ± 1.9% (Fig. 3D, Supplementary Table 9). Similarly, the severe patients had a mean total number of T-cell responses per patient of 5.6 ± 8.7 responses, while the mild patients had a mean of 3.3 ± 4.6 responses per patient (Fig. 3F, Supplementary Table 9). The frequency of total antigen-specific CD8^+^ T-cells and the number of T-cell responses per patient gradually declined over subsequent months, but remained higher for the severe patients compared to the mild patients throughout the investigated timespan (Fig. 3B-3E). Generally, mild patients showed a faster contraction of CD8^+^ T-cell responses than severe patients, with the most significant difference observed at TP3 (around 2 months after infection) (Fig. 3B-3E). However, the mean frequency of antigen-specific CD8⁺ T-cells decreased to similar levels, reaching 0.8 ± 0.9% in severe and 0.5 ± 0.5% in mild patients (Fig. 3D, Supplementary Table 9). Similarly, the mean number of T-cell responses decreased to 1.7 ± 2.6 in severe patients and 1.5 ± 1.7 in mild patients, respectively (Fig. 3E, Supplementary Table 9).

**Fig. 2.**
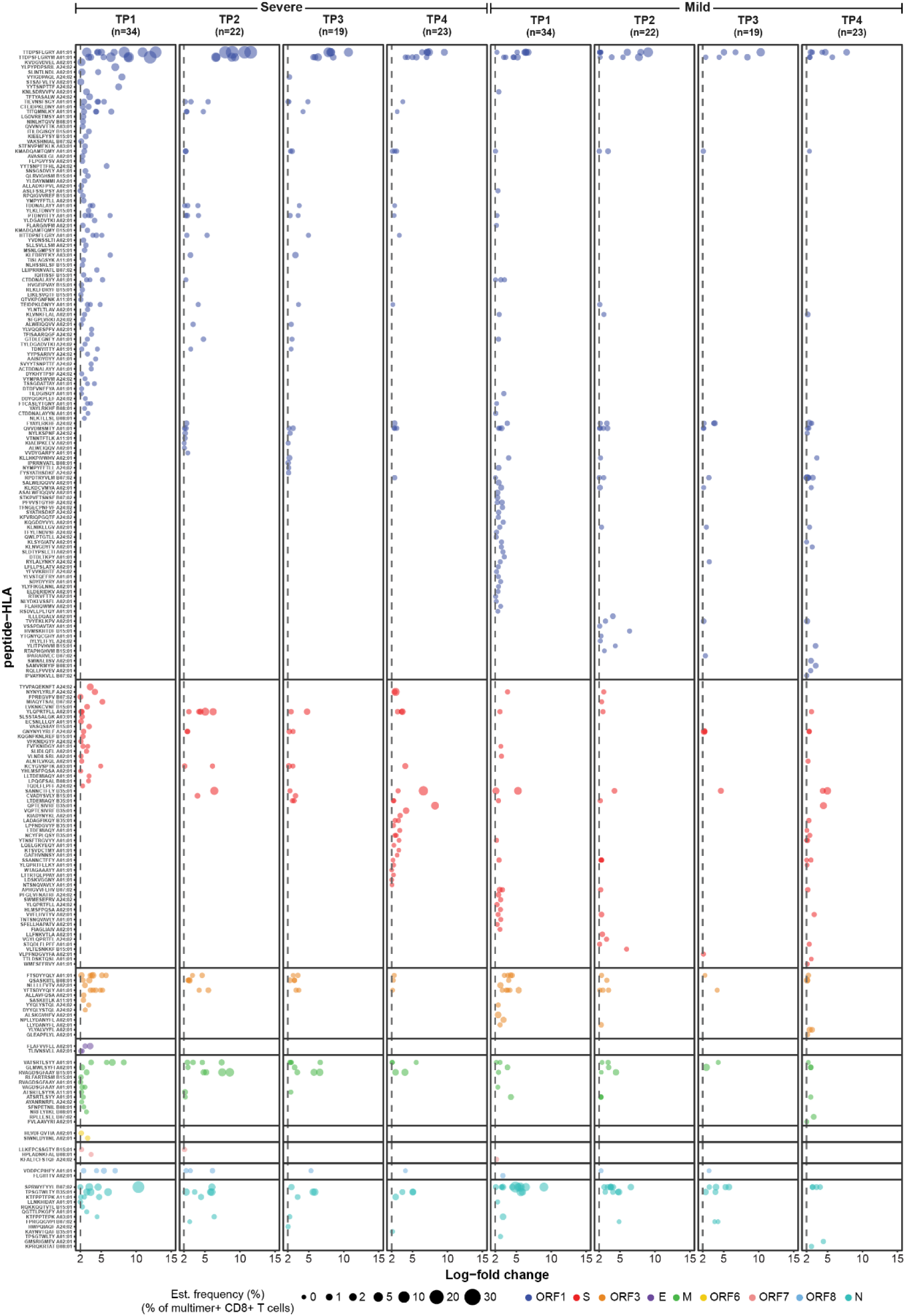
**Summary of SARS-CoV-2-specific T-cell responses in the severe and mild COVID-19 patients at four time points**. 601 pMHC were screened at TP1-TP4 and an additional 357 Spike-specific pMHC were included at TP4 for SARS-CoV-2 vaccinated patients. In parentheses is the number of patients analyzed for each time point and disease severity. Responses were identified based on the enrichment of DNA barcodes associated with each of the tested pMHC specificities (Log-fold change ≥ 2 and p < 0.001, analyzed using Barracoda). Only significant responses were included in the plot. Each dot represents one peptide-HLA combination per patient and is colored according to their protein of origin. The size of each significant response is proportional to the estimated frequency calculated from the percentage read count of the associated barcode out of the percentage of CD8^+^ multimer^+^ T-cells. SARS-CoV-2-specific T-cell responses were segregated based on their protein of origin.

**Fig. 3.**
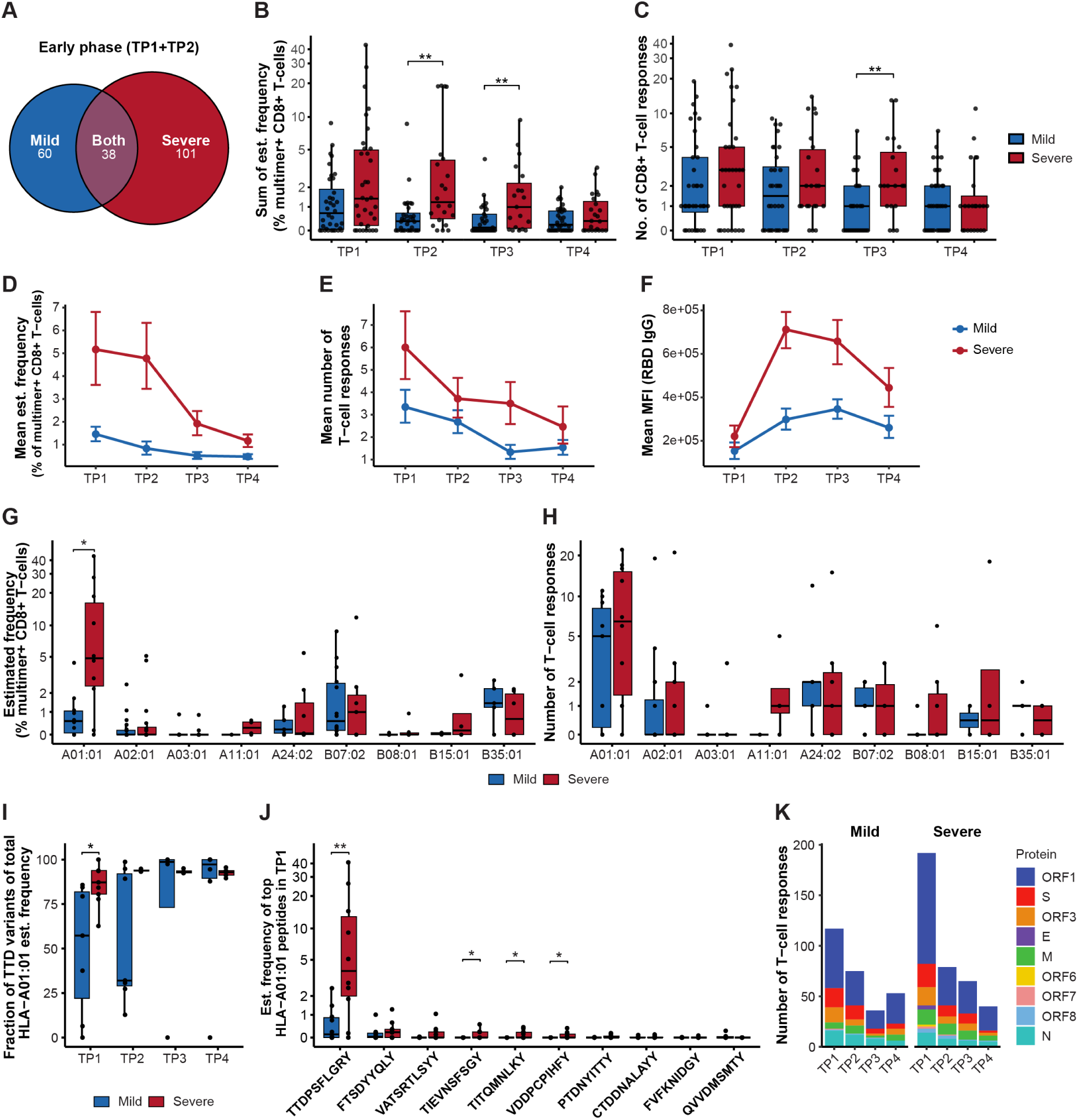
Kinetics of CD8^+^ T-cells and antibody levels in mild and severe COVID-19 patients. (**A**) Venn diagram shows the number of unique epitopes identified in mild, severe, or both groups of COVID-19 patients during the early phase post-diagnosis (TP1 + TP2). Box plots compare the sum of estimated frequency (%) (**B**) and the total number (**C**) of significant SARS-CoV-2-specific CD8^+^ T-cell responses between severe and mild COVID-19 patients across the four time points (TP1-TP4). Kinetics in the mean estimated frequency (%) (**D**) and the mean number (**E**) of SARS-CoV-2-specific T-cell responses in severe and mild patients across the four time points. (**F**) Kinetics of mean IgG antibody levels against the SARS-CoV-2 RBD across the four time points for severe and mild patients. Comparison of the sum of estimated frequency (%) (**G**) and the total number (**H**) of significant SARS-CoV-2-specific T-cell responses per HLA type at TP1 between severe and mild patients. (**I**) Bar plot compares the fraction of the sum of estimated frequency of the TTD variant peptides (TTDPSFLGRY, TTDPSFLGRYM and HTTDPSFLGRY) relative to the total HLA-A01:01-specific peptide responses in mild and severe patients. (**J**) Comparison of the estimated frequency of the top 10 HLA-A01:01-restricted peptides, selected based on prevalence, between mild and severe patients at TP1. TTDPSFLGRY and FTSDYYQLY also include the estimated frequency of their variant peptides (TTDPSFLGRYM and HTTDPSFLGRY for TTDPSFLGRY and YFTSDYYQLY for FTSDYYQLY). (**K**) Stacked bar plot summarizes the number of T-cell responses derived from each SARS-CoV-2 protein across the four time points in mild and severe patient cohorts. Statistical significance was calculated using Fisher’s exact test (K) and Mann-Whitney test (B, C, G, H, I, J). Significance levels are indicated as **** (p < 0.0001), *** (p < 0.001), ** (p < 0.01) and * (p ≤ 0.05).

We next determined the kinetics of T-cell responses in comparison to the humoral response (Fig. 1F). Plasma IgG levels were measured against Nucleocapsid (N), Spike receptor-binding domain (RBD), Spike 1 (S1), and Spike 2 (S2) antigens. TP4 samples of vaccinated individuals were excluded from the analysis to avoid the influence of vaccine-induced antibody levels. In contrast to the T-cell responses, which were detected at the maximum levels already at the first time point (TP1) (Fig. 3D, 3E), the antibody levels for both severe and mild COVID-19 patients remained low at TP1 and peaked only at the second time point (TP2) (Fig. 3F, Supplementary Fig. 4A). This delayed antibody response compared to the CD8^+^ T-cell response aligns with previous observations (23–25). Similar to the T-cell response, patients with severe COVID-19 consistently exhibited higher antibody levels across all time points (Fig. 3F, Supplementary Fig. 4A). A significant positive correlation between disease severity and both the peak levels and the duration of the antibody response has previously be described by (18,26,27). However, we did not observe a strong correlation between the T-cell frequency and the antibody levels, except at TP1 for Spike S1 antigen (R² = 0.34, p<0.001) in severe patients (Supplementary Fig. 4B).

Altogether, longitudinal analysis of T-cells showed a mutually exclusive antigen-specific T-cell repertoire in mild and severe disease and a strong association between the magnitude of CD8^+^ T-cell activation and severe COVID-19 in the acute phase of SARS-CoV-2 infection. However, the long-term memory pool of T-cell frequency and the breadth of T-cell repertoire contracts to a comparable level between mild and severe patients.

### Enhanced T-cell frequency and HLA-specificity strongly associated with COVID-19 disease severity

HLA polymorphism can strongly influence disease outcomes (28,29). Several studies across different population groups have shown protection or susceptibility of SARS-CoV-2 infection and disease outcomes associated with different HLAs (30–34). Since significantly higher frequencies of CD8^+^ T-cells were identified in severe COVID-19 patients, we next investigated their association with HLA and antigen specificity. Despite the equal distribution of HLA-A*01:01 in mild and severe patient groups (Supplementary Fig. 5A), HLA-A*01:01-specific T-cell frequencies were significantly higher in severe compared to mild patients, and these represent the highest T-cell frequencies observed across all HLA restriction investigated (Fig. 3G, Supplementary Fig. 5B). Statistical comparison across all the nine HLAs for the frequency of T-cells showed significant enrichment of HLA-A*01:01-specificities in the severe patient group (Mann-Whitney test, p = 0.015 (TP1); p = 0.006 (TP2), p = 0.012 (TP3), p = 0.005 (TP4)) (Fig. 3G, Supplementary Fig. 5B, Supplementary Table 11). Similarly, the number of HLA-A*01:01-restricted immunogenic epitopes was also higher for the severe patients (Mann-Whitney test, p = ns (TP1); p = 0.012 (TP2), p = 0.023 (TP3)), p = 0.009 (TP4)) (Fig. 3H, Supplementary Fig. 5C, Supplementary Table 11). It was also observed that the total percentage of estimated frequency and total number of HLA-A*01:01-restricted immunogenic epitopes (normalized to the number of patients per HLA) was highest compared to all other HLAs in the severe group (Supplementary Fig. 5D, 5E). Importantly, during the early phase of infection (TP1) in severe patients, the frequency of CD8^+^ T-cells specific to HLA-A*01:01-restricted antigens reached as high as 44% in individual patients (mean ± SD: 11.57 ± 14.37%). In contrast, the maximum frequency in mild patients was only 4.4% (mean ± SD: 0.95 ± 1.32%) (Fig. 3G, Supplementary Table 11). Among all HLA-A*01:01 responses, the ORF1-derived TTDPSFLGRY peptide (including its variants TTDPSFLGRYM and HTTDPSFLGRY) was the most dominant immunogenic epitope, eliciting the highest T-cell frequency in both mild and severe patients (Fig. 3I, 3J). However, the frequency of TTDPSFLGRY-specific T-cells was nearly tenfold higher in severe patients (mean ± SD: 10.31 ± 13.45%) than in mild patients (mean ± SD: 0.59 ± 0.84%) at TP1 (Fig. 3J).

Furthermore, we observed a higher prevalence of HLA-B*07:02 donors in the mild patient group (37.8%; 14 of 37 patients) compared to the severe patient group (19.4%; 7 of 36 donors) (Supplementary Table 6), suggesting a potential association between HLA-B*07:02 and mild COVID-19. This aligns with previous findings showing a strong association between the HLA-B*07:02 restricted SPRWYFYYL–specific T-cell response and mild disease, possibly due to cross-reactivity with the highly similar OC43/HKU-1-CoV N105–113 peptide (LPRWYFYYL) (31). Indeed, in our cohort, the frequency and prevalence of the most immunodominant HLA-B*07:02–restricted epitope (SPRWYFYYL; Nucleocapsid epitope, N105–113) were higher in mild patients, although the differences were not statistically significant (Supplementary Fig. 5F, Supplementary Tables 9 and 10). Finally, we compared the impact of different SARS-CoV-2 proteins on the T-cell response between mild and severe patients and found no difference in the number of T-cell responses across all time points. (Fig. 3K; Supplementary Table 13).

In summary, we identify a strong association of severe COVID-19 with HLA-A*01:01-restricted CD8^+^ T-cell response, dominated by a single antigen-specificity.

### COVID-19 vaccination broadens antigen-specific T-cell repertoire and boosts long-term memory

Since 17 patients (severe, n = 10; mild, n = 7) of our cohort received either one or two doses of a COVID-19 vaccine between TP3 and TP4 (Supplementary Table 2), we next evaluated the impact of vaccine-induced hybrid immunity. In both mild and severe patients, vaccination induced a significant increase in plasma IgG antibody levels against one or more of the vaccine-specific SARS-CoV-2 antigens (RBD, S1, and S2), indicating the vaccine’s efficacy in enhancing IgG antibody levels in patients across both severe and mild COVID-19 cases. However, post-vaccination IgG levels targeting Spike S1 and S2 were significantly higher in severe patients compared to mild patients (Fig. 4A and Supplementary Fig. 4C). This difference is likely due to the stronger infection-induced antibody responses in severe patients, which were maintained across all time points prior to vaccination (Supplementary Fig. 4A). Next, to assess the impact of vaccination on the T-cell repertoire, at TP4 we analyzed an additional 415 peptides (predicted to bind the nine HLAs included in this study) covering the entire SARS-CoV-2 Spike protein (GenBank ID: QHD43416.1), encoded by the Pfizer-BioNTech BNT162b2 mRNA vaccine (35) and the Moderna mRNA-1273 vaccine (36) (Supplementary Table 14). A notable increase in the frequency and number of Spike-specific T-cell responses was observed in vaccinated compared to non-vaccinated individuals at TP4 (Fig. 4B, 4C). Analysis of post-vaccination Spike-specific T-cell responses at TP4, compared to the earlier time points, identified T-cell responses to 14 different Spike-specific pMHC complexes across all vaccinated patients. Thus, SARS-CoV-2 vaccination further broadened the CD8^+^ T-cell repertoire either by boosting the frequency of existing infection-induced T-cells or generating de novo vaccine-specific responses that were not detected before vaccination. Additionally, post-vaccination analysis of the extended Spike library (tested only at TP4) identified T-cell reactivity to 12 additional Spike pMHC complexes across all vaccinated patients (Fig. 4D). Furthermore, even though the frequency of Spike-specific T-cells was significantly increased only in the severe patient group post-vaccination compared to pre-vaccination (Fig. 4E), both patient groups were efficient in enhancing existing responses or mounting new responses in response to vaccination (Fig. 4F). These results show that vaccination (post-infection) further boosts existing T-cell responses as well as broaden the overall CD8^+^ T-cell repertoire irrespective to the outcome of the natural infection.

**Fig. 4.**
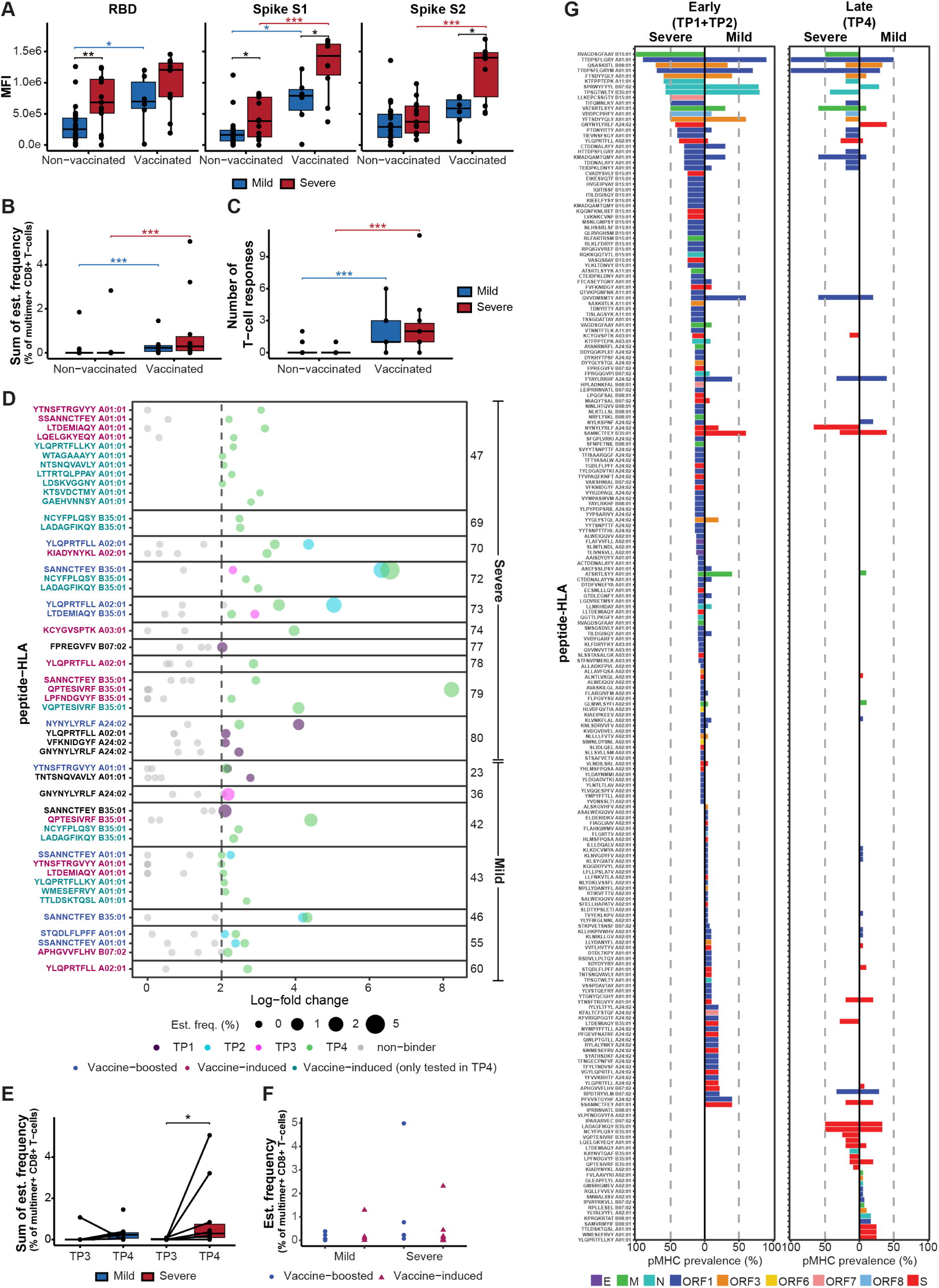
Analysis of SARS-CoV-2 antigen-specific T-cell immunodominance and vaccine-driven hybrid immunity in mild and severe COVID-19 patients. (**A**) Post-COVID-19 vaccination (TP4) levels of IgG antibodies against SARS-CoV-2 Spike protein subunits S1 and S2, and RBD in mild and severe COVID-19 patients. Box plot comparing the sum of the estimated frequencies (**B**) and the number (**C**) of SARS-CoV-2 Spike-specific T-cell responses at TP4 between mild and severe patients and between vaccinated and non-vaccinated groups. (**D**) Summary of SARS-CoV-2 Spike-specific T-cell responses across four time points in the 17 vaccinated patients. Significant responses to individual pMHC pairs are colored based on the time point at which they were identified and separated by Patient ID on the *y*-axis. Grey dots represent peptides without significant enrichment. Peptide-HLA labels on the *y*-axis are color-coded to indicate the type of response observed at TP4: blue, vaccine-boosted; pink, vaccine-induced; turquoise, vaccine-induced responses tested only at TP4; and black, T-cell responses detected at other time points but not at TP4. (**E**) Box plot comparing the sum of the estimated frequencies of SARS-CoV-2 Spike-specific T-cell responses between mild and severe vaccinated individuals pre-(TP3) and post-(TP4) vaccination. (**F**) Estimated frequency of SARS-CoV-2 Spike-specific T-cell responses identified post-vaccination in mild and severe COVID-19 patients. Only Spike pMHCs (n = 149 pMHC) analyzed during the complete longitudinal analysis are included in this plot. (**G**) Prevalence according to disease severity of CD8^+^ T-cell recognition towards SARS-CoV-2 epitopes detected in early (TP1+TP2) and late (TP4) time points. Late prevalence includes T-cell responses towards additional Spike peptides included only at T4. Only pMHC tested in more than 2 donors were included in this analysis. A dotted line is placed at 50% of prevalence to distinguish immunodominant epitopes. Bars are colored according to their protein of origin. (A, B, C, E) Mann-Whitney test, **** (p < 0.0001), *** (p < 0.001), ** (p < 0.01) and * (p ≤ 0.05).

### Immunodominant SARS-CoV-2 epitopes establish long-term CD8^+^ T-cell memory

Next, we evaluated the immunodominance and long-term prevalence of antigen-specific CD8 ^+^ T-cells. For robustness, only epitopes tested in at least three individuals with the corresponding HLA molecule per time point and severity group were included (Fig. 4G, Supplementary Fig. 6, and Supplementary Table 12). In the early phase following infection (TP1 and TP2), several highly prevalent epitopes (prevalence ≥ 50%) identified in severe patients were also observed in mild patients, while a significant number of epitopes detected in severe cases were not found in mild cases, and vice versa. Severe patients exhibited a broader repertoire of SARS-CoV-2-specific T-cell epitopes and a higher prevalence of T-cell recognition for individual epitopes. Among the severe patients, 8 epitopes were determined as immunodominant, based on T-cell recognition in more than 50% of the tested samples and detection in at least three or more patients. In mild patients, 7 immunodominant epitopes were identified. The epitopes HLA-A*01:01-TTDPSFLGRY (and its variant TTDPSFLGRYM), HLA-B*07:02-SPRWYFYYL, and HLA-B*35:01-TPSGTWLTY were considered immunodominant in both groups.

At TP4 (approximately 220 days post-infection), long-term memory was observed to have a more selective repertoire, primarily consisting of highly prevalent epitopes during the acute phase, along with the vaccine-boosted or vaccine-driven responses in vaccinated patients. Altogether, the long-term CD8^+^ T-cell memory was established against 43 and 47 epitopes in severe and mild patients respectively. The long-term immunodominance was found for six of these epitopes in the severe patient group with TTDPSFLGRY (and its variant TTDPSFLGRYM; HLA-A*01:01 restricted) being the most dominant response with 100% prevalence. In the mild patient group, none of the antigen-specific T-cells showed an immunodominant response (prevalence ≥ 50%), with TTDPSFLGRY-specific T-cells having a 50% prevalence in this group. Interestingly, the frequency of some of the most immunodominant CEF-specific T-cells establishing long-term memory pools was much higher as compared to the long-term frequency of the SARS-COV-2 antigen-specific T-cells (Supplementary Fig. 7A, 7B, Supplementary Table 15).

### Differential T-cell activation and memory profile in mild and severe COVID-19 patients

Next, we followed the phenotype kinetics of pMHC multimer-positive CD8^+^ T-cells based on the expression of cell surface markers (Fig. 1F, Fig. 5, Supplementary Fig. 2). SARS-CoV-2-specific T-cells from severe patients showed strong expression of CD38, HLA-DR, and PD-1 early in with gradual decline towards TP4. In mild cases, the expression of these activation markers was consistently lower. CD39 and CD69 levels were similar at early phase, however, a significantly higher expression of these markers was observed in mild patients at TP4 (Fig. 5A). Importantly, the differential activation profile between severe and mild COVID-19 patients was observed only for SARS-CoV-2-specific CD8⁺ T-cells, except for CD38 expression, which showed significant variation between mild and severe patients also among CEF-specific CD8⁺ T-cells (Supplementary Fig. 7C). Further, SARS-CoV-2-specific CD8⁺ T-cells exhibited significantly higher expression of activation markers compared to CEF-specific T-cells in both severe and mild patients, particularly during the early phase of infection (Supplementary Fig. 7D). Additionally, no variation was observed in the overall levels of CEF-specific multimer-positive T-cells between mild and severe patients or between the initial sampling at TP1 and the final sampling at TP4 (Supplementary Fig. 7A, 7B, Supplementary Table 15).

**Fig. 5.**
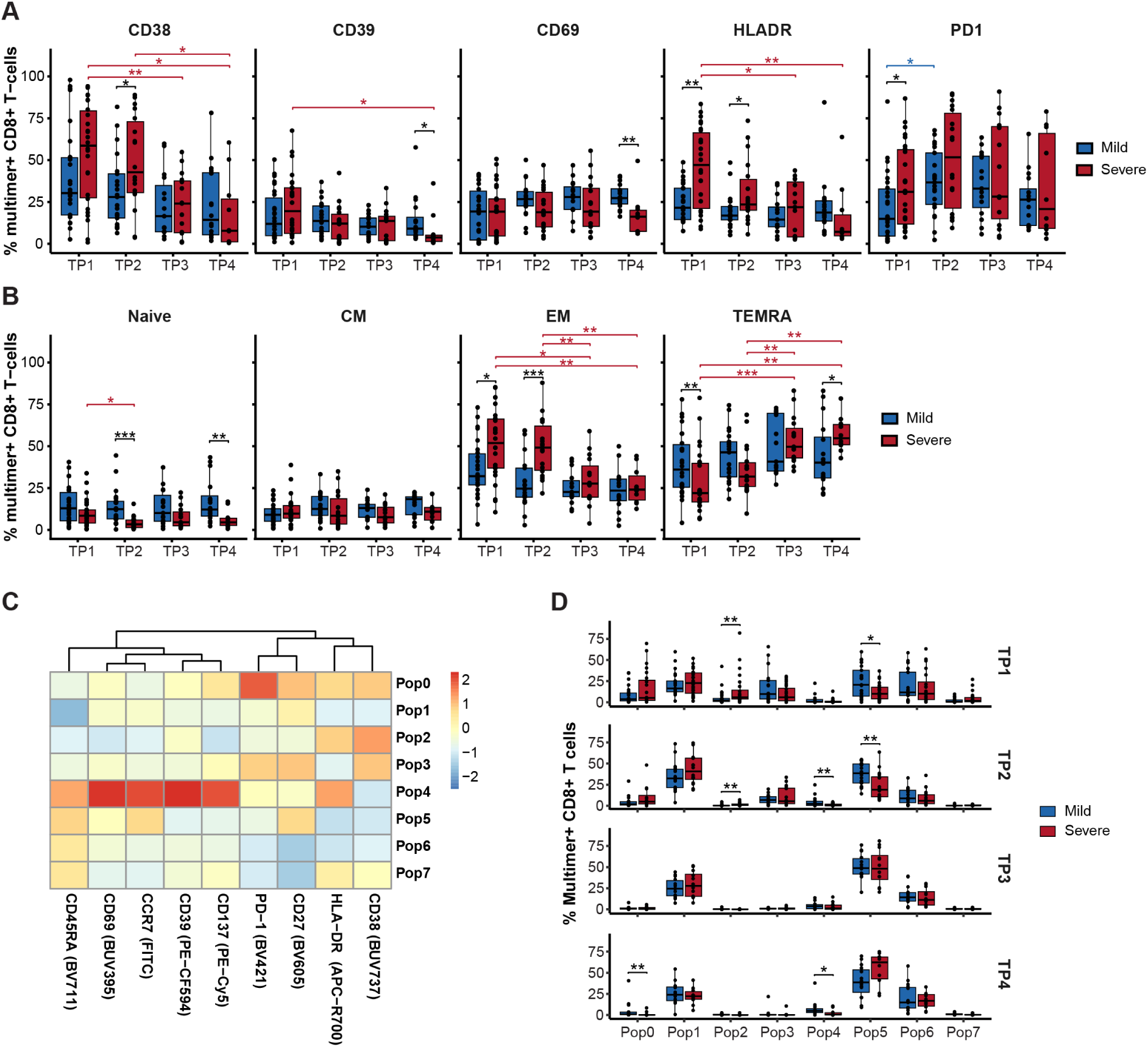
Memory and activation profile of SARS-CoV-2-specific T-cells in mild and severe COVID-19 patients. (**A**) Box plots indicating the percentage of pMHC multimer^+^ CD8^+^ T-cells expressing the surface markers (CD38, CD39, CD69, HLA-DR, and PD-1). (**B**) Memory phenotype (Naive, CM, EM and TEMRA) of pMHC multimer^+^ CD8^+^ T-cells based on the expression of CD45RA and CCR7 cell surface markers. (**C**) Heatmap based on mean fluorescence intensity (MFI) of each marker in each population identified by FlowSOM analysis (Supplementary Fig. 8A). (**D**) Box plots comparing the distribution of multimer positive CD8^+^ T-cells between mild and severe patients for each FlowSOM population across the different time points. (A, B, F) Mann-Whitney test between disease severity and Mann-Whitney test adjusting p-values with the Bonferroni method for comparison between time points, **** (p < 0.0001), *** (p < 0.001), ** (p < 0.01) and * (p ≤ 0.05).

Next, we compared the memory profile of SARS-CoV-2 multimer-positive CD8^+^ T-cells based on the expression of CD45RA^+^ and CCR7^+^ markers (37). The majority of the multimer^+^ memory CD8^+^ T-cells at all time points were either effector memory (EM; CD45RA^low^, CCR7^low^) or terminally differentiated effector memory cells re-expressing CD45RA (TEMRA; CD45RA^hi^, CCR7^low^), and low percentages of circulating naïve and central memory (CM) cells (Fig. 5B). Noticeably, at the acute phase of infection (TP1), antigen-specific T-cells of mild patients displayed a significant enrichment of the TEMRA subset compared to severe patients. In contrast, the acquisition of TEMRA profile in severe patients was slower and reached a significantly higher level than T-cells of the mild patients only at TP4. On the other hand, the EM subset of T-cells was significantly higher in severe COVID-19 patients at early time points (TP1 and TP2).

We further evaluated the phenotypic characteristics of SARS-CoV-2-specific CD8^+^ T-cells by performing dimensionality reduction with uniform manifold approximation projection (UMAP) (38) and unsupervised clustering of multimer^+^ CD8^+^ T-cells using the FlowSOM algorithm (39). UMAP showed distinct distributions of the SARS-CoV-2 multimer-reactive T-cells between severe and mild patients, highlighting different expression profiles of T-cell memory and activation markers (Supplementary Fig. 8A). A heatmap generated by FlowSOM displayed the relative mean fluorescence intensity (MFI) for each marker across each clustered population (Fig. 5C). This unbiased clustering analyses confirms the findings from the manual gating, namely enrichment of populations with EM phenotype and activation markers in servere patients, demonstrated by significant enrichment of populations 2 at TP1 and TP2. On the contrary for mild disease patients, significant enrichment of population 5 was observed, representing a cell populations with TEMRA/naïve phenotype and minimal activation signature (Fig. 5C, 5D, Supplementary Fig. 8B).

Altogether, our longitudinal data of SARS-CoV-2-specific CD8^+^ T-cells in mild patients show an early onset of a TEMRA-like effector profile with an overall lower expression of activation markers, whereas severe patients had a strong EM profile with enhanced expression of activation markers. This data suggest that mild patients have stronger recruitment of memory T-cells compared to severe patients.

### Single-cell analysis reveals enhanced cytotoxic T-cell profile in mild COVID-19 patients

To evaluate the functional consequences of the substantially high frequency of antigen-specific T-cells observed in severe patients, we performed single-cell analysis of antigen-specific T-cells from six COVID-19 patients (mild, n = 3; severe, n = 3) at early (TP1 or TP2) and late (TP4) time points post-infection (Fig. 6A). We selected six immunogenic peptides (HLA-A*01:01 restricted TTDPSFLGRY, FTSDYYQLY, VATSRTLSYY, and CTDDNALAYY; HLA-B*07:02 restricted SPRWYFYYL; HLA-A*02:01 restricted YLQPRTFLL) identified in our pMHC multimer analysis of these patients at both early and late time points (Supplementary Table 10 and Supplementary Table 16). PBMCs were incubated with either individual peptides or an HLA-matching pool of peptides, and cells were sorted for single-cell analysis based on the expression of T-cell activation markers CD69 and CD137 (Supplementary Fig. 9 and Supplementary Table 16). Our single-cell analysis combined transcriptomic profiling, oligo-tagged antibody staining (CITE-seq), and T-cell receptor sequencing (TCR-seq) (Fig. 6A). After demultiplexing and quality control assessment, 2784 cells were classified as singlets and used for downstream analysis (Supplementary Fig. 10 and 11).

**Fig. 6.**
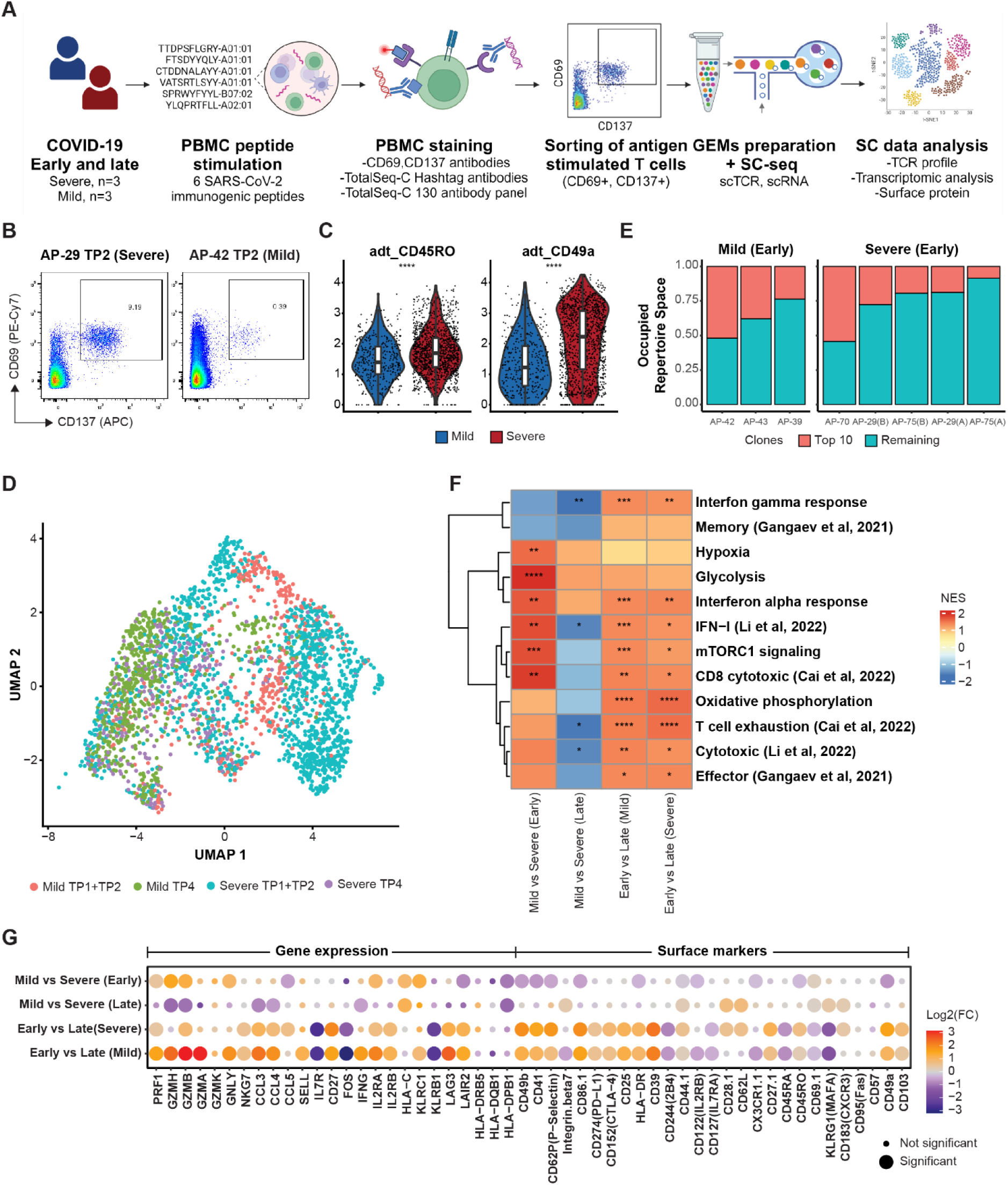
Single-cell analysis comparing antigen-specific CD8^+^ T-cells of mild and severe COVID-19 patients. (**A**) Experimental pipeline for single-cell transcriptome, surface proteome, and TCR analysis of three mild and three severe patients analyzed during the early and late phase of COVID-19 infection. (**B**) Representative flow cytometry plots showing antigen-stimulated CD8^+^ T-cells sorted for single-cell analysis based on CD69 and CD137 expression. (**C**) Comparison of surface markers expression levels between severe and mild samples for early time point. Mann-Whitney test: *p < 0.05, **p < 0.01, ***p < 0.001, ****p < 0.0001. (**D**) UMAP plot of scRNA-seq data colored by a combination of severity and time point. (**E**) TCR clonal composition within early (TP1/TP2) samples for mild and severe conditions. Patient ID labels (*x*-axis) are described in Supplementary Table 16. TCR clones are merged according to proportion into two categories: the top 10 represented clones and the remaining ones. (**F**) Gene set enrichment analysis (GSEA) for selected gene sets. Heatmap of normalized enrichment score (NES) coupled with adjusted p-values (*p < 0.05, **p < 0.01, ***p < 0.001, ****p < 0.0001). (**G**) Selected genes and surface markers (columns) identified from differential expression analysis between cells grouped by severity and time point (rows). Significant defined by adjusted p-value < 0.05.

Since the pMHC multimer-based flow cytometry analysis showed early onset of memory T-cell with TEMRA and EM profile in mild and severe patients respectively (Fig. 5), we compared the T-cell phenotype of mild and severe patients using single-cell CITE-seq analysis. At early time points, T-cells of severe patients showed higher expression of cell surface markers associated with antigen-specific memory markers (CD44, CD122, CX3CR1, CD45RO), and tissue-resident marker CD49a, whereas T-cells of the mild patients showed enhanced expression of costimulatory molecule CD28 and the resident memory marker CD69 (Fig. 6C, Supplementary Fig. 12A). On the contrary, at the late time point (about six months post-infection), T-cells of mild patients showed higher expression of activated memory T-cells; CD44, CD62L, CD28, and KLRG1 (Supplementary Fig. 12B).

Next, we looked at the UMAP distribution of T-cells based on the COVID-19 disease severity across early and late time points. Cells from severe and mild patients clustered differently at early time points, however at the late time point clusters were more homogenous for the two disease types, suggesting that COVID-19 disease severity is influence by T-cell activation at early stage of infection (Fig. 6D). We did not observe any differences in T-cell receptor (TCR) clonal distribution as well as total TCR-repertoire space occupied by larger clones (compared top 10 TCR clones from each sample) among the mild and severe patients (Fig. 6E). Interestingly, gene set enrichment analysis of multiple pathways (40–43) related to T-cell activation and function showed that at early time point T-cells of the mild patients displayed significantly higher activation signatures than T-cells of the severe patients, for pathways such as CD8^+^ cytotoxicity and interferon (IFN) response. In addition, pathways for mTORC1 signaling, glycolysis, and hypoxia were also significantly activated, potentially suggesting a better activation of cytotoxic CD8^+^ T-cells in mild patients compared to severe patients during the early onset of the infection (44–47). At the late time point these differences were diminished and showed comparable or more activation profile for the T-cells of the severe patients. Furthermore, comparing early and late time points of both severe and mild patients showed significantly higher activation in the early phase of the infection across many of these pathways (Fig. 6F and Supplementary Fig. 13).

To gain gene-specific insight at the transcriptome and proteome level of the higher cytotoxic profile identified in T-cells of the mild patients, we performed differential gene expression analysis on a single-cell level and pseudobulk level (Supplementary Fig. 13). In agreement with the observation made from the gene set enrichment analysis, we found significantly elevated expression of several genes linked to T-cell effector and cytotoxic function such as perforin (*PRF1*), granzymes (*GZMH, GZMB*), and granulysin (*GNLY*) in T-cells of the mild patient at early time points. Contrary to this, these genes’ expression was higher in severe patients when compared at the late time point (Fig. 6G). Furthermore, the increased cytotoxic profile of T-cells from mild patients was significantly dominated at the early time points compared to the late time point which was less significant for severe patients (comparing early and late TPs). The cell surface expression of CD49b, CD41, CD39, and CD69 supports the activation profile observed at the transcriptomic level for T-cell activation at early time points in both mild and severe patients, however, fails to capture the dynamics of differential activation of T-cells in mild vs severe patients (Fig. 6G).

In conclusion, our single-cell data identifies lower functional activation and reduced cytotoxic capacity of CD8^+^ T-cells from severe patients in the acute phase of SARS-CoV-2 infection.

### Immunodominant antigen-specific T-cells are differentially activated in mild and severe COVID-19 patients

With the observations of a substantially higher magnitude of T-cell activation in severe COVID-19 patients (Fig. 3) and the reduced cytotoxicity of such T-cells, as identified from the single-cell analysis (Fig. 6), we next examined the association of antigen-specificity and differential functional profile in mild and severe disease. The strong correlation between the frequency of HLA-A*01:01-restricted SARS-CoV-2-specific T-cells and severe COVID-19 is mostly due to the extremely high frequency of TTDPSFLGRY (TTD)-specific T-cells, and TTD is also the most immunodominant antigen and establishes long-term memory in both severe and mild patients (Fig. 3J, Fig. 4G, Fig. 7A, Supplementary Table 12). Immunodominance of TTD antigen is SARS-CoV-2 infection has also been reported previously by us and others (14,42,48). Thus, we focused our analysis on TTD-specific T-cells.

**Fig. 7.**
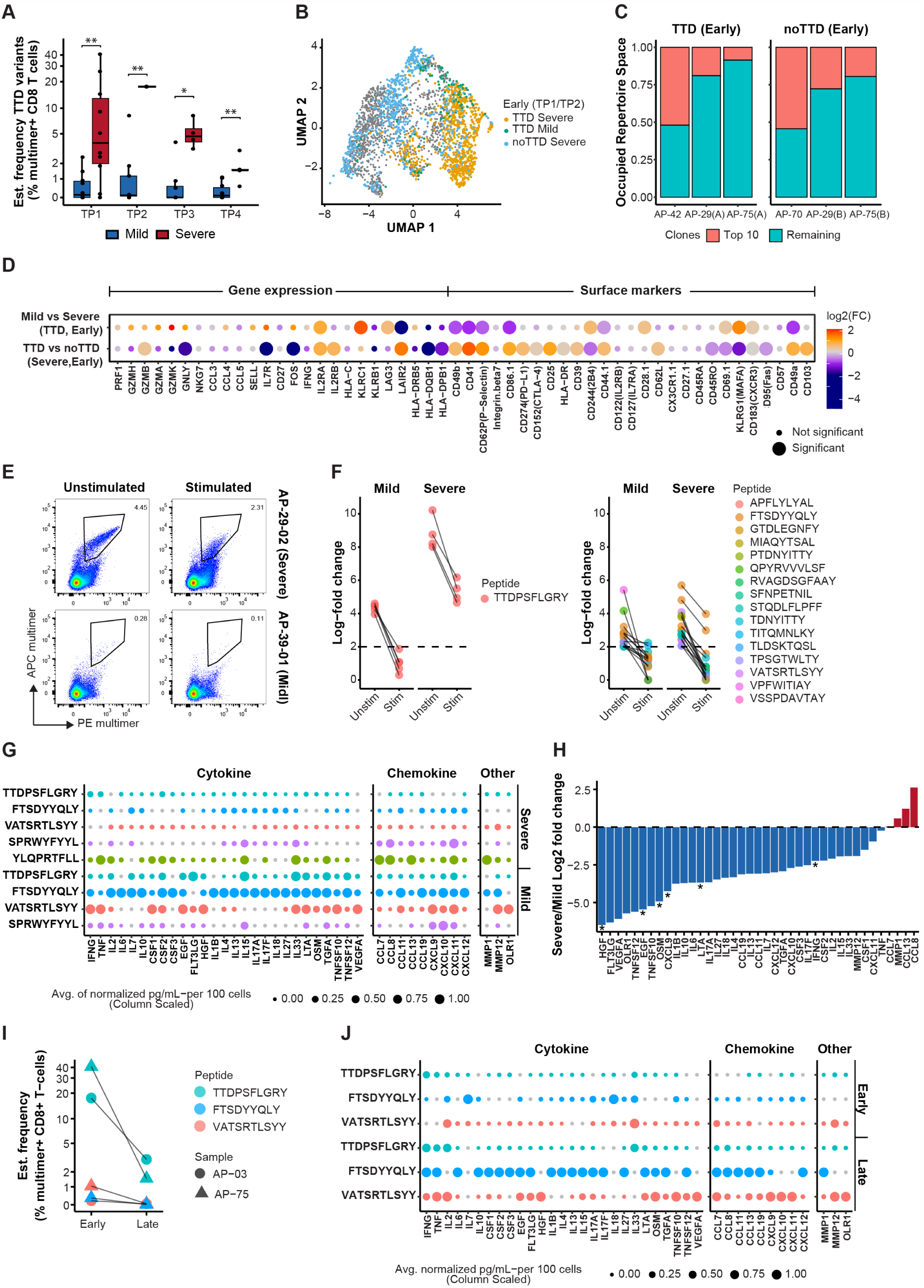
Functional assessment of antigen-specific T-cells in mild and severe COVID-19 patients. (**A**) Sum of estimated frequencies of HLA-A01:01-TTD (and its variants; TTDPSFLGRYM, and HTTDPSFLGRY)-specific T-cells in mild and severe COVID-19 patients across the different time points. Mann-Whitney test, mild vs severe; TP1 (p = 0.00685), TP2 (p = 0.00685), TP3 (p = 0.0107), TP4 (p = 0.00733). P-values were adjusted using the Bonferroni method and found non-significant when comparing across time points. (**B**) UMAP plot of scRNA-seq data for early (TP1/TP2) time points colored by a combination of peptide stimulation groups (TTD and no TTD) and disease severity. (**C**) Clonal composition of TTD-and no TTD-specific T-cell responses within early-phase (TP1/TP2) samples. Clones are grouped according to proportion into two categories: the top 10 most represented clones and the remaining clones. (**D**) Differential expression analysis of selected genes and surface markers (columns) between cells grouped by disease severity and peptide specificity (rows). Genes and markers with significant differences (adjusted p < 0.05) are indicated. (**E**) Representative flow cytometry plots of double-positive (PE^+^APC^+^) multimer-binding T-cell populations with and without peptide stimulation used to assess TCR down-regulation. Unstimulated controls were treated with equimolar DMSO. (**F**) Antigen-specific T-cells identified using pMHC multimers in peptide-stimulated samples compared to unstimulated controls (TCR down-regulation assay) in mild and severe patients. Only the significant pMHC responses (Log-fold change ≥ 2 and p < 0.001) identified in unstimulated controls were compared with the respective peptide-stimulated samples. **(G)** Mean protein secretion levels (pg/mL per 100 cells) of antigen-specific T-cells at the early phase (TP1/TP2) in severe (n = 3) and mild (n = 3) COVID-19 patients. Values were scaled to a range of 0 to 1. (**H**) Severe-to-mild ratio of mean protein secretion levels, calculated using normalized protein concentrations across all peptides for each condition. Mann–Whitney test, *p < 0.05. (**I**) Estimated frequency (%) of TTD-specific T-cells in severe patients at early and late phases post-diagnosis. **(J)** Magnitude of mean protein secretion levels (pg/mL per 100 cells) for each peptide, calculated as the average for all samples within each phase (early and late), following peptide stimulation of severe patient samples shown in panel I. Values were scaled to a range of 0 to 1. In (C, D), TTD refers to TTDPSFLGRY-specific T-cells, and no TTD refers to T-cells specific to other epitopes (FTSDYYQLY, VATSRTLSYY, CTDDNALAYY, YLQPRTFLL, or SPRWYFYYL).

In the single-cell analysis of antigen-stimulated cells, during the early phase of infection (TP1/TP2) TTD-specific T-cells of severe patients not only clustered separately from mild patients but also from the T-cells of other antigen-specificities (no TTD) (Fig. 7B). This suggests a differential activation state of TTD-antigen driven T-cells in the acute phase of infection. We evaluated if TCR clonotypes distribution was driving the observed differential activation of TTD-specific T-cells. However, no difference was observed at clonotype level on total clonal composition or enrichment of specific clones between TTD-specific and other peptide-specific T-cells (Fig. 7C). Next, comparing gene expression profiles based on differential gene expression analysis, TTD-specific T-cells of severe patients showed lower expression of cytotoxic molecules (*PRF1, GNLY, GZMH, GZMA, GZMB,* and *GZMK*) compared to mild patients but were not statistically significant (Fig. 7D). Similarly, within severe patients, compared to TTD-specific cells, T-cells specific to other antigens (no TTD) were more cytotoxic with higher expression of several genes related to T-cell cytotoxicity (*PRF1, GZMH, GZMA*, and *GZMK*) including a significantly higher expression of *GNLY*. On the contrary, *GNZB, IL2RA*, and *UL2RB* expression were higher in TTD-specific T-cells of severe patients, and the gene enrichment analysis further showed a significant increase in genes associated with interferon activation pathways of TTD-specific cells of severe patients compared to the mild (Fig. 7D, Supplementary Fig. 14A). Furthermore, TTD-specific T-cells of mild patients showed a significant increase in genes for *IL2, LAG3*, and *KLRC1*. On the cell surface level, TTD of mild patients showed significantly higher expression of CD69, KLRG1, CD28, and CXCR3. Importantly, the TTD-specific cells showed significantly up-regulated expression of cell surface markers associated with tissue localization and trafficking such as CD49a, CD49b, CD103, CD62 (P and L), and CD44. A higher expression of these markers combined with lower expression of KLRG1 and increased expression CXCR3 and of inhibitory markers (CTLA-4 and CD39) suggest a tissue-specific localization of TTD-specific CD8^+^ T-cells (Fig. 7D, Supplementary Fig. 14B) (49). Gene enrichment analysis further revealed up-regulated pathways related to glycolysis and hypoxia in T-cells specific to TTD in mild patients compared to severe patients (Supplementary Fig. 14A) and in severe patients the same was identified for T-cells not-specific for TTD (Supplementary Fig. 14B), suggesting potential metabolic deregulation of antigen-specific T-cells in severe COVID-19 cases. Since TTD-specific T-cells constitute the majority of antigen-specific T-cell populations in HLA-A*01:01– positive patients with severe COVID-19 disease (Fig. 3I), the lower activation profile of T-cells observed in these patients (Fig. 6F, 6G) compared to mild patients could be attributed to the immunodominance of these antigen-specific T-cells. The TTD antigen-specific metabolic deregulation is further supported by enhanced expression of the *UCP2* gene only in TTD-specific T-cells of severe patients (Supplementary Fig. 13D and 14B). *UCP2* is known for antigen-induced expression that reduces glycolysis and fatty acid synthesis to control T-cell differentiation (50). Next, to understand the role of TCR composition, we evaluated antigen-specific TCR gene segments for any preferential enrichment. We observed a higher V9-2 and V27 gene segment usage for alpha and beta chains respectively in TTD-specific cells compared to no TTD cells (Supplementary Fig. 15). Furthermore, an antigen-specific combination of different gene segments was evident in both alpha and beta chains across the evaluated T-cell specificities (Supplementary Fig. 16). Together, these data highlight possible dysfunction of the most immunodominant antigen-specific T-cells in severe patients.

### T-cells of severe COVID-19 patients show impaired TCR stimulation and cytokine secretion in the acute phase of the infection

To further investigate the findings of impaired antigen-specific T-cells in severe COVID-19, we used a recently established TCR down-regulation assay to probe the efficiency of T-cells upon peptide stimulation (51). Antigen-induced T-cell activation leads to TCR down-regulation and thus reduces the availability of cell-surface TCRs for pMHC multimer-based T-cell detection (51). Utilizing this approach, we used DNA-barcoded pMHC multimers to measure T-cell activation by comparing the level of TCR-down regulation (inversely proportional to the pMHC multimer binding) with and without antigen stimulation. 17 antigen-specific T-cell responses (including TTD) were studied in PBMCs collected at the early phase of infection (TP1 and TP2) from ten patients (including two severe and two mild patient samples studied in the single-cell analysis). After 24-hour peptide stimulation we observed almost complete disruption of pMHC multimer binding, resulting from TCR internalization, in samples in mild patients (n = 6). In contrast, in severe patients (n = 4), a substantial fraction of pMHC multimers binding capacity was maintained after peptide stimulation, suggesting a diminished functional response to the antigen stimuli (Fig. 7E, Supplementary Fig. 17). Since our pMHC multimers of each antigen-specificity were tagged with a DNA barcode, we compared the log2FC of individual antigen-specificity with and without peptide stimulation to access antigen-specific TCR-down regulation as a proxy to T-cell activation. Interestingly, TTD-specific T-cells of all four severe patients showed incomplete TCR-down regulation (log2FC >2, above the threshold of T-cell detection) in response to peptide stimulation. In contrast, TTD-specific T-cells of all the mild patients, as well as T-cells specific to other antigen specificities, showed complete TCR internalization (Fig. 7F). In addition to TTD-specific T-cells, we also identified another antigen-specific T-cell (FTSDYYQLY, HLA-A*01:01 restricted) in two out of the three severe patients with incomplete TCR internalization (Fig. 7F, right panel). Thus, our TCR down-regulation assay demonstrates functional impairment of SARS-CoV2-specific T-cells exclusively in severe patients.

Next, to quantify the functional impairment of T-cells in severe patients, we measured the amplitude of cytokine and chemokine production of CD8^+^ T-cells after antigen-specific stimulation using a 48-plex Olink Target Cytokine panel (Supplementary Table 18). We analyzed samples from six patients (mild, n = 3; severe, n = 3) collected at the early phase post-infection (TP1/TP2). Antigen-specific T-cells (including TTD) of severe patients secreted substantially lower amounts of cytokines and chemokines as compared to the T-cells of mild patients (Fig. 7G). Specifically, T-cells of mild patients showed overall higher secretion of most of the analytes including the CD8^+^ T-cell effector cytokines such as IFN-γ, TNF-α, IL-2, and IL-6 (Fig. 7H). To further evaluate if this dysfunctional state of antigen-specific T-cells in severe patients persists in the long-term memory pool, we evaluated early and late time point samples of two severe patients with the highest frequency of SARS-CoV-2-specific T-cells (Fig. 7I). For all three antigen-specificities T-cells from the late time point (six months post-infection) showed up to eightfold higher secretion of the cytokines and chemokines compared to the T-cells of acute phase of infection (Fig. 7J, Supplementary Fig. 18).

These data corroborate our findings of differential phenotype, transcriptomics, and metabolic activity of T-cells in mild and severe patients, and the dysfunctional state of CD8^+^ T-cells in the early phase of infection in severe patients.

## Discussion

Since the beginning of the COVID-19 pandemic, several studies have shown the important role of CD8^+^ T-cells in mounting an early and effective immune response (3,7,9,52,53). However, antigen-specific dynamics of differential T-cell activation associated with COVID-19 disease severity and their long-term impact are not fully resolved. In this study, we perform a comprehensive longitudinal profiling of CD8^+^ T-cells for 601 pMHC specificities for their association with COVID-19 disease. Our analysis of antigen-specific T-cell frequency, phenotype, and epitope-specific immunodominance provides valuable insight into antigen-specific differences in T-cell activation in mild and severe COVID-19 patients.

Our data identifies several critical features of CD8^+^ T-cell kinetics in SARS-CoV-2 infection that are strongly associated with COVID-19 disease. In the acute phase of infection, SARS-CoV-2-reactive CD8^+^ T-cell repertoire is driven by a large set of immunogenic epitopes. We identified 210 SARS-CoV-2 epitopes across the four time points in this patient cohort. Only 45 of these epitopes overlapped in mild and severe disease, and the remaining were unique to mild or severe patient groups. Contrary to the acute phase, the long-term CD8^+^ T-cell memory (based on the TP4 data) is established against only a small fraction of T-cell epitopes. Immunodominant epitopes with high prevalence (HLA-specific) in the acute phase contributed the most in establishing the long-term T-cell population, likely due to the higher initial frequency. In line with existing reports, COVID-19 vaccination boosted the T-cell frequency of the memory T-cells induced by primary infection (54–56). Importantly, vaccination induced existing T-cells and generated de novo responses in severe and mild patients, thus broadening the overall T-cell repertoire reactive to Spike and non-Spike epitopes. Although we have not tested the T-cell immunity concerning the evolving landscape of SARS-CoV-2 variants (57), the long-term memory constituted largely by epitopes from non-Spike regions of SARS-CoV-2 is likely to provide better protection as compared to T-cells specific to only Spike epitopes against different variants with high mutation frequency in Spike protein (58).

Previous studies have reported antigen-and HLA-specific association in asymptomatic or mild SARS-CoV-2 infection (31). Our data demonstrates a substantially high frequency of CD8^+^ T-cells and the number of T-cell responses in severe patients is strongly driven by HLA-A*01:01-restricted antigens. In the early outbreak of the COVID-19 pandemic in Italy and later in different geographical locations, HLA-A*01:01 populations have been found susceptible to severe COVID-19 (32–34). Persistent antigen stimulation has been shown to impair cytotoxic T-cell function in viral infections and cancer (59). In SARS-CoV-2 this has been largely observed in severe patients and could be due to prolonged viral encounters and antigen stimulation, also due to associated comorbidities and age (60,61). We identify such functional impairment in severe patients linked largely to a single epitope-specificity.

In this study, ORF1-derived HLA-A*01:01-restricted TTD antigen was the most immunodominant epitope in both mild and severe patients, however, the T-cell frequency was significantly higher in severe cases and constituted the majority of the total SARS-CoV-2 antigen-specific T-cells in individual patients. For example, in a specific severe patient, the total SARS-CoV-2 antigen-specific T-cell frequency reached as high as 40% of the total CD8^+^ T-cell population of which 33% were specific to TTD (and its homologs TTDPSFLGRYM and HTTDPSFLGRY). Our previous study showed reduced IFN-γ and TNF-α secretion capacity of CD8^+^ T-cells reactive to TTD epitope in severe patients, but not in mild patients (14). Similarly, another study also highlighted impaired cytokine secretion of TTD-specific cells in severe patients (42). By comparing multiple antigen-specificities in single-cell analysis and functional assessment, we now show that in severe patients functional impairment is largely driven by immunodominant epitope-specific T-cells that show altered metabolic reprogramming associated with T-cell exhaustion and reduced cytokine secretion capacity. To our knowledge, such large-scale expansion of antigen-specific T-cells has been observed in CMV infection (62) and a comparative assessment of antigen-specific T-cells from SARS-CoV-2 and CMV infection in the acute phase would be relevant to gain further understanding of antigen-specific dynamics of T-cell activation and impairment. Together with the high frequency of HLA-A*01:01-restricted CD8^+^ T-cells in severe patients and their dysfunctional state identified in the acute phase of the infection, our data strongly argue for CD8^+^ T-cell-associated disease severity.

However, it is important to emphasize that the early dysfunction of T-cells in severe patients seems to resolve over time as at the late time point T-cells of severe patients including TTD-reactive T-cells have comparable levels of transcriptomic markers and cytokine profile in response to peptide stimulation. Together, we hypothesize that early functional impairment of SARS-CoV2-specific CD8^+^ T-cells leads to a lack of viral control and consequent massive clonal proliferation to counteract the persistent viral load. Pending viral control the SARS-CoV2-specific CD8^+^ T-cell compartment are gradually reverting to their functional state. As such severe disease patients respond to vaccines at an equal level as patients with mild disease.

Furthermore, an increased magnitude of both the T-cell and antibody response was detected in severe patients, but compared to the antibody response, CD8^+^ T-cell response was established and peaked early in the acute phase of infection in both mild and severe patients. Early activation of cytotoxic T-cells in SARS-CoV-2 infection and vaccination has been shown previously and supports the importance of T-cells in early viral control (63–65). In contrast to the higher proliferation in severe patients, mild patients showed early recruitment of memory T-cells with TEMRA-like features. Existing data dissecting immune response in mild and asymptomatic SARS-CoV-2 infection support early activation of cytotoxic T-cells for effective viral clearance (12,13). T-cells specific to HLA-B*07:02-specific immunodominant epitope SPRWYFYYL were most prevalent in mild. We and others have shown the presence of HLA-B*07:02-SPRWYFYYL T-cells in SARS-CoV-2 unexposed individuals and sequence homology between SPRWYFYYL peptides and peptides derived from other human coronavirus (HCoVs) (14,66) and it has been associated with mild cases (31). Thus, a pre-existing T-cell memory might be involved in rapid expansion and effective viral control in mild or asymptomatic patients (3,48,67,68). Indeed, our flow cytometry data support this notion, as we observe an early activation and TEMRA profile and mild patients compared to severe patients, and we also detect a higher frequency of HLA-B*07:02-SPRWYFYYL-specific T-cells in mild patients. In the future, the identification of individual T-cell epitopes that induce early T-cell activation could be useful in determining immune memory as well as using such antigens for vaccine design (69).Our study has potential limitations. The stratification of patients into mild and severe categories is largely based on hospitalization and non-hospitalization and there is a large variation in hospitalization days (1 day to 36 days). Furthermore, our severe patient group also constituted patients with comorbidities which may influence COVID-19 disease severely and bias in T-cell immune response (70,71). Also, due to the limited cell number, our single-cell data lacked the strength to follow individual TCR clones from the early phase to the memory phase for individual antigen-specificity to investigate clonal dynamics of CD8^+^ T-cells in correlation to their functional characteristics (72). Lastly, for the 18 patients who were vaccinated after infection between TP3 and TP4, we evaluated the complete library of Spike-specific peptides only at TP4 and not at the earlier time points, thus, limiting the longitudinal analysis of some of the vaccine-induced T-cells across all time points in comparison to infection-induced T-cells.

In summary, by analyzing a large repertoire of SARS-CoV-2 immunogenic epitopes, we provide a comprehensive assessment of CD8^+^ T-cells in mild and severe COVID-19 patients for their contribution in the early phase of infection as well as in establishing long-term memory. We demonstrate the association of antigen-specific T-cell frequency and COVID-19 disease severity and show the potential functional impairment of antigen-specific T-cells in severe COVID-19 patients. In contrast, by combining flow cytometry and single-cell analysis, our data indicate an enhanced T-cell activation and memory profile in mild COVID-19 patients. Thus, demonstrating the utility of longitudinal profiling of T-cells on individual antigen level to understand clinical features concerning immune response that may have a significant impact in assessing T-cell immunity for therapeutic purposes, such as vaccine design, as well as in guiding future strategies involving novel pathogens.

## Materials and Methods

### Study design

This longitudinal study was designed to investigate the CD8^+^ T-cell mediated immune response against SARS-CoV-2 infection, and the influence of antigen-specific T-cells in severe and mild COVID-19 disease. We analyzed serial samples from 73 individuals with confirmed SARS-CoV-2 infection who presented mild-to-moderate and severe disease. To study T-cell recognition, immunodominance, long-term memory, and the phenotype of reactive T-cells, our approach integrated DNA-barcoded peptide-MHC (pMHC) multimers together with a 12-color flow cytometry panel. To further analyze the immune response, a single-cell analysis was conducted to provide detailed insights into the functionality, transcriptomic profiles, and TCR repertoires of the CD8^+^ T-cells activated during infection. Additionally, our study extended to analyze the T-cell response in 17 SARS-CoV-2 infected patients who were later vaccinated with a COVID-19 mRNA vaccine.

To complement our analysis of cellular immunity, we employed multiplex bead-based assays to profile antibody levels in plasma. This enabled us to correlate humoral responses with cellular immunity offering a comprehensive view of the immune response to SARS-CoV-2 infection.

### Clinical sample collection and ethical approval

The study protocol, including procedures for sample collection, was approved by the Committee on Health Research Ethics in the Capital Region of Denmark (H-20026375). The patient cohort comprised 73 individuals diagnosed with COVID-19 during the first wave of pandemic. Patients were stratified based on disease severity using an operational classification scheme developed specifically for this study, as no standardized criteria were established at the time. Stratification was based on clinical observations and hospital care requirements, dividing patients into four groups: 1 = non-hospitalized (mild symptoms), 2 = hospitalized with mild symptoms, 3 = hospitalized with severe symptoms, 4 = hospitalized requiring intensive care unit (ICU) support. All patients who required hospitalization (groups 2, 3, and 4) were classified as severe, except for two patients who were hospitalized due to pre-existing conditions and exhibited no COVID-19 symptoms; these two individuals were categorized as mild. The cohort included 36 patients classified as severe (median age = 62.4 years; mean hospitalization duration = 8 ± 7.17 days) and 37 patients classified as mild (median age = 47.2 years). Participants were included if they were 18 years or older, had a COVID-19 diagnosis confirmed by RT-PCR testing, and provided written informed consent. Detailed demographic and clinical characteristics of the participants, including age, gender, disease symptoms, disease severity, and comorbidities are provided in Supplementary Table 1.

Blood samples were systematically collected at four time points. The initial collection (TP1) occurred within two weeks (9±7 days) post-diagnosis. The subsequent collections were scheduled at 27±10 days (TP2), 58±10 days (TP3), and 218±30 days (TP4) after diagnosis. Of the cohort, 17 patients received one or two doses of a COVID-19 mRNA vaccine (either Pfizer BioNTech BNT162b2 or Moderna mRNA-1273) between TP3 and TP4. (Fig. 1A, Supplementary Fig. 1 and Supplementary Table 2).

Peripheral blood mononuclear cells (PBMCs) were isolated from the collected blood samples by density gradient centrifugation using Leucosep tubes (Greiner Bio-One 227288) with Lymphoprep media (StemCell Technologies 07861). Following isolation, PBMCs were cryopreserved at −150°C in FCS (Gibco) + 10% DMSO, while the plasma portion was stored at-80°C for future analysis. Additionally, PBMC samples underwent genotyping for HLA-A, B, and C loci utilizing next-generation sequencing techniques (DKMS Life Science Lab GmbH, Germany) (Supplementary Table 6).

### SARS-CoV-2 peptide selection

For this study, we selected 553 HLA class I-binding peptides across 9 prevalent HLA alleles, covering 9 proteins from the SARS-CoV-2 isolate Wuhan-Hu-1 (GenBank ID: MN908947.3), previously identified as immunogenic. Of these, 311 epitopes were identified in our previous study (Saini *et al*., 2021 (14)). An additional 33 peptides were included based on genome-wide T-cell mapping performed in 19 COVID-19 patients at TP1 using DNA barcoded pMHC multimers. In this mapping, 2198 peptides were predicted to bind at least one of seven selected HLA alleles (HLA-A*01:01,-A*02:01,-A*03:01,-A*24:02,-B*07:02,-B*08:01, and –B*15:01) using NetMHCpan-4.1 with a rank threshold of <1% (21), and subsequently screened for CD8^+^ T-cell reactivity (Supplementary Table 4). The remaining peptides were selected from IEDB (www.iedb.org, Immune Epitope Database (16)). based on immunogenic epitopes reported by other studies and restricted to the HLA alleles included in this study with a rank threshold of <1% predicted using NetMHCpan-4.1 (21). The 553 peptides were used to construct 601 peptide-HLA pairs for experimental assessment (Supplementary Table 3).

To assess T-cell reactivity in samples from COVID-19 vaccinated individuals, 278 additional Spike peptides (357 pMHC pairs). These peptides were predicted using NetMHCpan-4.1 (21) to bind one or more of the nine prevalent HLA-A and HLA-B molecules, applying a binding rank threshold of <1% (Supplementary Table 14). This resulted in a comprehensive set of 415 peptides spanning the complete SARS-CoV-2 Spike protein encoded by the Pfizer BioNTech BNT162b2 mRNA vaccine and the Moderna mRNA-1273 vaccine (GenBank ID: QHD43416.1). This expanded set was used to generate 506 Spike peptide-HLA pairs for experimental evaluation at TP4.

Furthermore, 38 peptides from cytomegalovirus (CMV), Epstein-Barr virus (EBV), and influenza (FLU), collectively referred to as CEF peptides, were incorporated to enable comparative analyses with SARS-CoV-2-derived peptide-reactive T-cells (Supplementary Table 8). All peptides were custom-synthesized by Pepscan (Pepscan Presto BV, Lelystad, The Netherlands), dissolved in 10 mM DMSO, and stored at-20°C until required for experimental procedures.

### MHC class I monomer production

The production of all nine types of MHC-I monomers followed established protocols (73,74). In summary, recombinant MHC-I heavy chains and human β2-microglobulin (β2m) were produced as inclusion bodies in *Escherichia coli* employing pET series expression vectors. The expressed proteins were refolded with the aid of UV-sensitive peptide ligands specific to each HLA, facilitating the correct assembly of the molecules (74). HLA-A*02:01 and A*24:02 variants were refolded and purified empty as previously described (75). Following the refolding process, the MHC-I molecules were biotinylated utilizing the BirA biotin-protein ligase reaction kit (Avidity LLC, Aurora), and then purified using size exclusion chromatography (SEC-HPLC; Waters Corporation, USA). The monomers were stored at-80°C for future applications.

### Generation of DNA-barcoded multimer libraries

The generation of DNA-barcoded multimer libraries for SARS-CoV-2-and CEF-derived peptides followed the protocol established by Bentzen *et al.* (76). Individual peptide–MHC (pMHC) complexes were formed by incubating each peptide with its respective MHC molecule. For HLA-A*02:01 and A*24:02, direct peptide loading was employed (75), while UV-mediated peptide exchange was used for the remaining HLA types (74). The pMHC monomers were then coupled to allophycocyanin (APC) or phycoerythrin (PE) using dextran molecules tagged with unique DNA barcodes.

### Antigen-specific T-cell identification

Patient HLA-matching SARS-CoV-2 and CEF pMHC multimer libraries were pooled and incubated with PBMCs for 15 minutes at 37°C, as described previously (76). This step was followed by a 30-minute incubation at 4°C with a panel of phenotype antibodies and a dead cell marker to assess cell viability (Supplementary Table 7). After staining, cells were fixed in 1% paraformaldehyde for preservation.

Cells were analyzed using a FACSAria flow cytometer (AriaFusion, BD Biosciences), where pMHC multimer-binding CD8^+^ T-cells were identified and sorted (Supplementary Fig. 2).

### Analysis of TCR down-regulation upon antigen stimulation

Ten patient samples (mild, n = 6; severe, n = 4) collected during the early phase post-infection (TP1/TP2) were analyzed for activation-induced TCR down-regulation upon SARS-CoV-2 antigen stimulation, as described by (51). PBMCs from the samples were resuspended in X-Vivo media (Lonza BE02-060Q) supplemented with 5% human serum and stimulated using a SARS-CoV-2 peptide pool including 40 HLA-A*01:01-, 15 HLA-A*02:01-, 15 HLA-B*08:01-, and 15 HLA-B*35:01-restricted peptides (1 μM of each peptide) (Supplementary Table 17) or with DMSO (concentration-matched negative control) for 24 hours at 37°C. Following stimulation, cells were washed to remove peptides or DMSO and stained using a pool of DNA barcode-labeled multimers conjugated with PE and APC, prepared as previously described, along with a lineage antibody phenotype panel and a dead cell marker (Supplementary Table 7). DNA barcode-labeled multimers were generated for the 100 specificities used for stimulation (Supplementary Table 17). All double-positive (PE^+^ and APC^+^) pMHC multimer-binding CD8^+^ T-cells were analyzed and sorted using a FACSAria flow cytometer (AriaFusion, BD Biosciences) (Supplementary Fig. 17).

### DNA barcode sequence analysis

DNA barcodes from sorted multimer positive CD8^+^ T-cell populations, as well as from an aliquot of the multimer pool (to serve as a baseline), were amplified using the Taq PCR Master Mix Kit (Qiagen, 201443) together with sample-specific forward primers to enable accurate sample identification. The amplification products were purified using the QIAquick PCR Purification Kit (Qiagen, 28104) and sequenced at PrimBio (USA) using an Ion Torrent PGM 316, 318, or an Ion S5 530 chip (Life Technologies).

Sequencing data derived from DNA barcodes were analyzed using the Barracoda software package (76) (https://github.com/SRHgroup/Barracoda-2.0). This tool was employed to determine the number of sequencing reads and clonally reduced reads associated with each pMHC-specific DNA barcode. It also calculated the fold changes (FC) in read counts for each sample compared to the average counts from triplicate baseline samples, along with the p-values and false discovery rates (FDRs) as detailed in (76). Criteria for significant T-cell responses were established based on DNA barcodes having an FDR < 0.1% (equivalent to p < 0.001) and a Log2FC>2 when compared to baseline values across the entire pMHC library. The frequency of T-cells for each significantly enriched barcode was calculated from its percentage read count relative to the total CD8^+^ multimer^+^ T-cell population. To ensure specificity in T-cell response detection, a non-HLA-matching, non-SARS-CoV-2 infected healthy donor was included as a negative control. Peptides identified in this control sample were excluded from the analysis.

In our analysis of infection-specific CD8^+^ T-cell responses, we excluded Spike-specific responses that demonstrated an increased frequency at TP4 compared to TP3 in patients who were vaccinated between TP3 and TP4. This approach was taken to remove the impact of vaccine-induced responses at TP4 from our evaluation.

### Flow cytometry analysis

Flow cytometry data were analyzed using FlowJo data analysis software (version 10.9.0; FlowJo LLC). The phenotype analysis of SARS-CoV-2 and CEF pMHC multimer positive CD8^+^ T-cells was conducted according to the gating strategy shown in Supplementary Fig. 2. FCS files of samples were concatenated at both the APC-positive and negative-population gates (500 events for each per sample). Concatenated files were visualized using Uniform Manifold Approximation and Projection (UMAP, Version 3.3.3, FlowJo plugin (38) and clustered by FlowSOM (FlowJo plugin (39)) analysis based on the markers CD38, CD39, CD69, CD137, HLA-DR, PD-1, CCR7, CD45RA, and CD27. A heatmap of the FlowSOM metaclusters was automatically generated by the plugin to display the relative expression of each marker within each metacluster. The heatmap was adjusted to show the expression in each cluster relative to the mean fluorescence intensity (MFI) values of each marker, instead of relative to the MFI values of all markers.

### Quantification of anti-SARS-CoV-2 IgG antibodies using multiplex immunoassay

The multiplex bead-binding assay Bio-Plex Pro Human IgG SARs-CoV-2 N/RBD/S1/S2 4-Plex Panel (Bio-Rad, 12014634) was used to quantify anti-SARS-CoV-2 IgG antibodies specific for Nucleocapsid (N), Spike receptor-binding domain (RBD), Spike 1 (S1) and Spike 2 (S2) antigens. Plasma samples were centrifuged at 1000xg for 10 minutes before being diluted at a 1:100 ratio using the sample diluent provided by the kit and processed as per manufacturer’s instructions for analysis. The assay included both a pre-mixed positive control, comprising human IgG antibodies against the four targeted SARS-CoV-2 antigens, and a negative control to validate the assay’s specificity and sensitivity. The sample diluent also served as a blank to measure background non-specific binding levels. All samples and controls were analyzed in single measurements. Median fluorescence intensities (MFI) were captured using the Bio-Plex MAGPIX Multiplex Reader (Bio-Rad Laboratories), operated with the xPONENT software version 4.2 (Luminex Corporation). To account for dilution and non-specific background signals, the raw MFI values were adjusted by multiplying by the dilution factor and then subtracting the background MFI.

### Data processing and statistics

T-cell recognition data, determined by DNA-barcoded pMHC multimers analysis and Barracoda software, was plotted using R Studio (77). The ggplot2 package version 3.4.4 (78) was used to generate box, bar, column, scatter and dot plots for data visualization and the venn diagrams were generated using the eulerr package version 7.0.0 (79). Dot plots for visualization of T-cell responses displays peptide sequences with no significant enrichments as gray dots and all peptides with a negative enrichment are set to LogFC equal zero. Significant responses were colored based on different criteria for each plot. Fig. 2 only shows significant responses. The size of each significant response is proportional to the estimated frequency (%) calculated from the percentage read count of the associated barcode out of the percentage of CD8^+^ multimer^+^ T-cells. Box plots were generated for data quantification and visualization, and their related statistical analyses were performed using package rstatix version 0.7.2 (80). Mann-Whitney test was used for unpaired comparisons and in cases where the test was used to compare samples across different time points the p-values were adjusted using the Bonferroni method to correct for multiple comparisons. For statistical analysis of Fig. 3K Fisher’s exact test with Hochberg-Benjamini multiple testing correction (significance level: p<0.05) was conducted using package stats version 4.3.2 (77). The correlation coefficient (r^2^) and p-values shown in Supplementary Fig. 4B were generated using a linear model.

### T-cell staining and sorting for single-cell analysis

Six COVID-19 patients (severe, n = 3; mild, n = 3), identified to have T-cell response through DNA barcode-labeled pMHC multimers were selected for single-cell analysis. For each patient a sample collected early after SARS-CoV-2 infection (TP1 or TP2) and a sample collected at a late time point (TP4) were included for analysis. PBMCs from the selected samples were cultured in X-vivo media + 5% HS and stimulated with 1 µM of SARS-CoV-2 peptides for 24 hours at 37°C, 5% CO_2_. Refer to Supplementary Table 16 for details on peptides used for each sample. Following peptide stimulation, cells were washed with Cell Staining Buffer (PBS + 0.5% BSA) and resuspended in 20 µL of the buffer. Five µL of Human TruStain FcX Fc Blocking reagent was added to the cells, and incubated 10 minutes at 4°C. The TotalSeq-C Human Universal Cocktail (BioLegend 399905) phenotype panel was reconstituted with 27.5 µl of cell staining buffer. Subsequently, 12 µl of the reconstituted cocktail were added to each sample and incubated for 15 minutes at 4°C. Then, 0.5 µl of hashing antibodies (BioLegend, TotalSeq-C anti-human Hashtag 1-16 Antibodies, Supplementary Table 16), an antibody solution mix and a dead cell marker (Supplementary Table 7) were added to each sample and incubated for 30 minutes at 4°C. Cells were washed three times in cell staining buffer and maintained on ice until acquisition. Activated CD8^+^ T-cells were sorted based on the activation markers CD69 and CD137 using a FACS Melody (BD) (gating strategy provided in Supplementary Fig. 9). Approximately 20,000 cells from all samples were collected into a single tube containing 100 μL of Cell Staining Buffer, centrifuged at 390 g for 10 min at 4°C, and the supernatant discarded.

### Construction of single-cell libraries

The preparation of Gel Beads-in-emulsion (GEMs) and downstream processing of DNA barcodes and mRNA was performed utilizing the 10x Genomics 5’ v2 chemistry according to the manufacturer’s protocol (Chromium Next GEM Single Cell 5’ Reagent Kits v2 (Dual Index), with the Feature Barcode technology for Cell Surface Protein & Immune Receptor Mapping) (10x Genomics, USA). Sorted cells were loaded into a single lane of a Chromium Next Generation Chip and run in a Chromium Controller (10x Genomics, USA) to generate individual GEMs, followed by synthesis of cDNA from poly-adenylated mRNA, and DNA from cell surface protein Feature Barcode derived from antibodies. After 16 cycles of targeted amplification, the products were separated according to size using SPRIselect beads (Beckman Coulter, B23318), and processed separately for the construction of TCR (VDJ), 5ʹ Gene Expression (GEX), and Barcode (BC) libraries. The libraries were quantified with the Qubit dsDNA HS Assay Kit (Invitrogen Q32851) and combined at a ratio of 1 BC: 5 GEX: 1.5 TCR. Libraries were sequenced at Novogene Company Limited (UK) on a NovaSeq system (Ilumina) running a 150 paired-end program.

### Single-cell RNA and ADT analysis

Raw FASTQ files were aligned to the human reference genome GRCh38 v1.2.0 and V(D)J reference v.5.0 with Cell Ranger v.7.1.0 (81) using command cellranger multi with default parameters. Reference for the feature layer was constructed according to the barcodes that have been used in the experiment. The resulting matrices were loaded as a Seurat object (Seurat v5.0.2 (82)) and analyzed. Demultiplexing was performed using HTODemux with standard parameters after Centered Log-Ratio (CLR) normalization for hashing antibodies with more than 10.000 read counts. Next, singlets were filtered to exclude cells with more than 2000 or less than 200 expressed genes with ≥ 5% detected mitochondrial genes. Only protein coding (biomaRt v2.58.2 (83)) and mitochondrial genes were considered for gene expression assay. We used LogNormalize method for gene expression assay and CLR normalization for antibody derived tag (ADT) assay.

After scaling, 2000 most variable genes were selected for the PCA, and 18 principal components were used for clustering and UMAP visualization. We noticed that the first principal component is correlated with nFeature RNA metrics and affects further steps and clustering, thus, we concluded that resulting clusters are more likely related to technical variation in our data rather than biological one and we focused on a comparison between conditions (mild vs severe patients and early vs late time points) instead of cluster-based analysis. Also, a small cluster of cells was found with high expression of heat shock proteins and excluded for further analysis (cluster 5, Supplementary Fig. 11D).

Gene expression markers for the resulting clusters were calculated using FindAllMarkers with minimal percentage = 0.25 and logFC threshold = 0.25. Differentially expressed genes between conditions were calculated on a cell basis (FindMarkers, minimal percentage = 0, logFC threshold = 0) or using pseudobulk approach (AggregateExpression and DESeq2 v1.42.1 (84)). Gene set enrichment analysis (40) was applied to ranked statistics after differential expression on cell level (-log10(p_val)*sign(avg_log2FC)) using fgsea package (85) and gene sets from MSigBD (hallmark gene sets (40)) and selected publications (Li *et al.*, 2022 (41), Gangaev *et al.* 2021 (42), Cai *et al.* 2022 (43)). We used ComplexHeatmap (86) and EnhancedVolcano (87) packages for the visualization of differential expression analysis results.

### TCR profiling data analysis

Package scRepertoire (88) was used to analyze the filtered contig annotations. We called clones “strict”-VDJC gene and CDR3 nucleotide and set the filterMulti as TRUE, allowing us to select the top 2 represented chains for cells with multiple chains.

### Single-cell Statistics

Mann–Whitney test was used to compare two independent groups. Analyses were performed in R Studio version 4.3.3 (77).

### Functional characterization of T-cells using the Olink platform

Six patient samples (mild, n = 3; severe, n = 3) collected at early phase post-infection (TP1/TP2), along with two additional samples from patients with severe disease collected at a late phase (TP4), were analyzed using proximity extension assay technology (Olink Proteomics AB, Sweden). PBMCs (1 × 10⁶ cells) were cultured in X-vivo media supplemented with 5% HS and incubated for 24 hours at 37°C. After incubation, cells were washed three times with PBS to remove residual HS, resuspended in X-vivo media, and stimulated with either 1 μM of individual SARS-CoV-2 epitopes or DMSO (concentration-matched negative control) for an additional 24 hours at 37°C (Supplementary Table 18). Following stimulation, cell supernatants were collected for analysis. The DTU Centre for Diagnostics (Denmark) conducted the analysis using the Olink® Target 48 Cytokine panel (Olink Proteomics AB, Sweden), which enables simultaneous quantification of 45 inflammatory protein biomarkers. Quality control was ensured by running the Olink® Target 48 Sample Control in triplicate. Each sample was analyzed in a single measurement, and protein concentrations were reported in pg/mL.

For data analysis, protein concentrations from DMSO-stimulated samples were subtracted from those of SARS-CoV-2 peptide-stimulated samples to determine the specific response to viral epitopes. Measurements exceeding the upper limit of quantification (ULOQ) were set to the ULOQ value for the respective protein, while those with NaN readings were adjusted to 0 pg/mL. Normalization was performed based on the proportion of peptide-specific T-cells in each sample to determine the expected protein concentration per 100 cells (Supplementary Table 10).

Data visualization was performed using the ggplot2 package (version 3.4.4) (78) in R Studio. For dot plot visualizations, numerical data were row-scaled to a range of 0 to 1 to facilitate comparison and interpretation (Fig. 7G, 7J). Scaling was performed using the rescale function from the scales package (version 1.3.0) in R Studio. Differences between severe and mild disease, as well as early and late conditions, were analyzed by calculating fold changes for each analyte. Fold changes were derived from the mean pg/mL, calculated using the normalized protein concentrations of all peptides under each condition. Statistical comparisons were conducted using the Mann–Whitney test (Fig. 7H and Supplementary Fig. 18).

## Acknowledgments

We thank all patients for participating and contributing the analyzed samples; S. Sebbaha, A. G. Burkal T. M. L. Nguyen, B. Rotbøl, A. F. Løye for excellent technical support, and all the collaborators for active participation to this work.

## Author contributions

S.P.A.H. designed and performed experiments, analyzed data, prepared figures, and wrote the manuscript. K.K.M. and S.M.S.-G. analyzed the data and prepared figures. K.D. analyzed the data, prepared figures and wrote the manuscript. M.K. and T.T. performed experiments. D.S.H. designed the research, facilitated patient samples and clinical data collection. A.O.G. conceived the idea, supervised clinical study, patient participation, clinical data, and sample collection. S.R.H. conceived the idea, designed, and supervised the study. S.K.S. conceived the idea, designed and supervised the study, analyzed the data, and wrote the manuscript.

