## Supplementary Material for "Comprehensive longitudinal profiling of SARS-CoV-2-specific CD8^+^ T-cells reveal strong functional impairment and recognition bias as markers for disease severity"

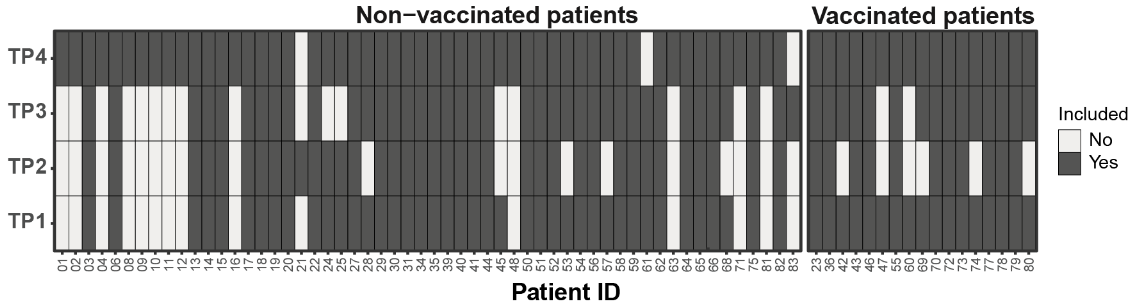


**Supplementary Fig. 1. Clinical samples details.** PBMC samples included per time point for each COVID-19 patient.

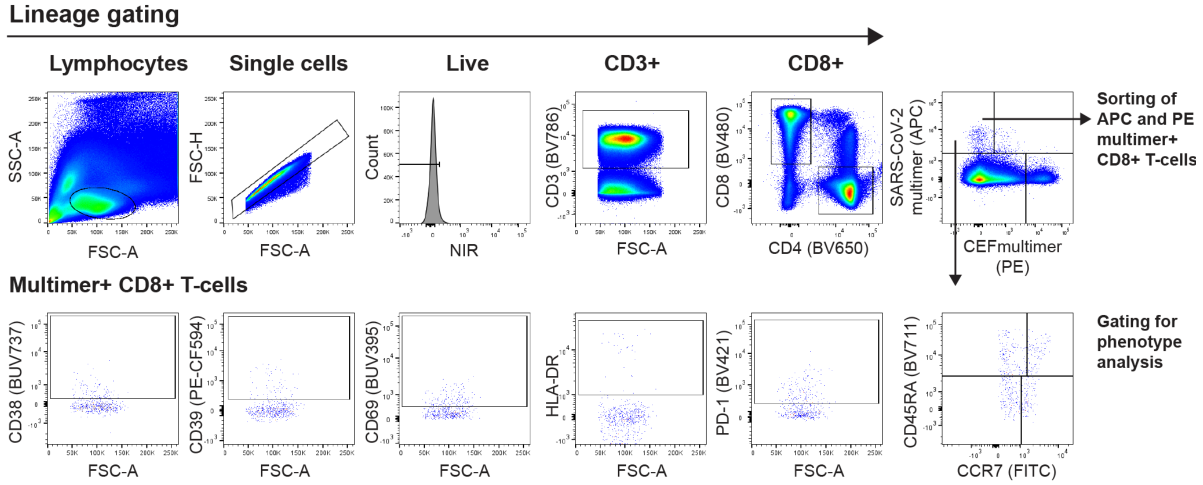


**Supplementary Fig. 2. Flow cytometry gating strategy for sorting and phenotyping SARS-CoV-2 multimer$^{+}$ CD8$^{+}$ T-cells.** Representative flow cytometry plots illustrating the gating strategy applied to PBMCs from COVID-19 patients. PBMCs were stained with DNA-barcoded pMHC multimers and surface antibody markers to sort SARS-CoV-2 (APC) and CEF (PE) multimer$^{+}$ CD8$^{+}$ T-cells and to quantify multimer$^{+}$ CD8$^{+}$ T-cells expressing phenotype markers.

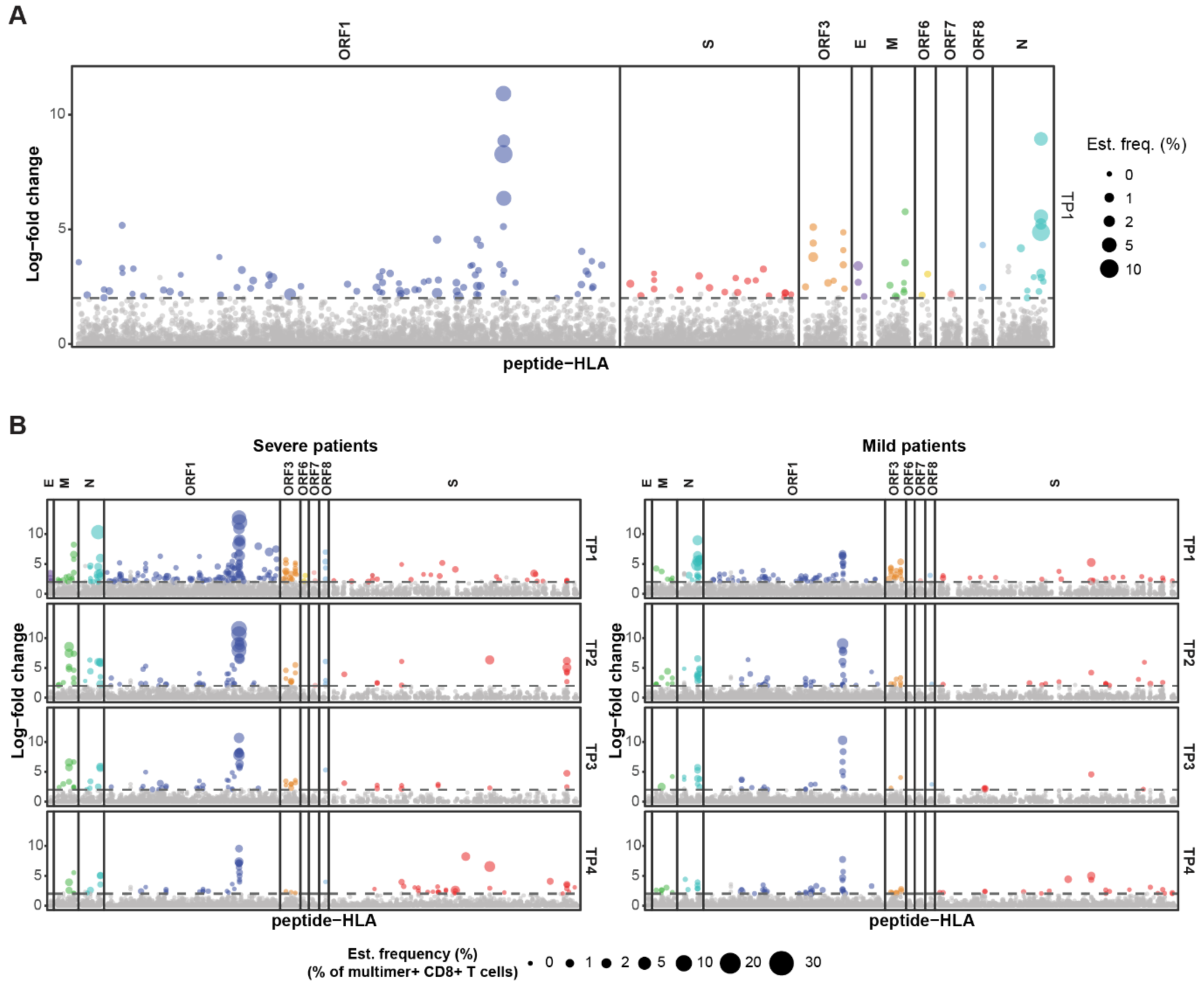


**Supplementary Fig. 3. Summary of SARS-CoV-2-specific $CD8^+$ T-cell responses in COVID-19 patients.** (**A**) $CD8^+$ T-cell recognition to individual peptides across COVID-19 patients (n = 19) at TP1, identified during epitope mapping of the complete SARS-CoV-2 genome (Supplementary Table 4). (**B**) Longitudinal profiling of SARS-CoV-2-specific T-cell responses across four time points (TP1–TP4) in COVID-19 patients, categorized by protein of origin and disease severity. (A, B) Individual epitopes were identified based on the enrichment of DNA barcodes associated with specific peptide-HLA combinations (Log-fold change>2 and $p < 0.001$, analyzed using Barracoda). Each dot represents one peptide-HLA combination per sample. Significant responses are colored according to their protein of origin and their size is proportional to the estimated frequency (%) calculated from the percentage read count of the associated barcode out of the percentage of $CD8^+$ multimer$^+$ T-cells.

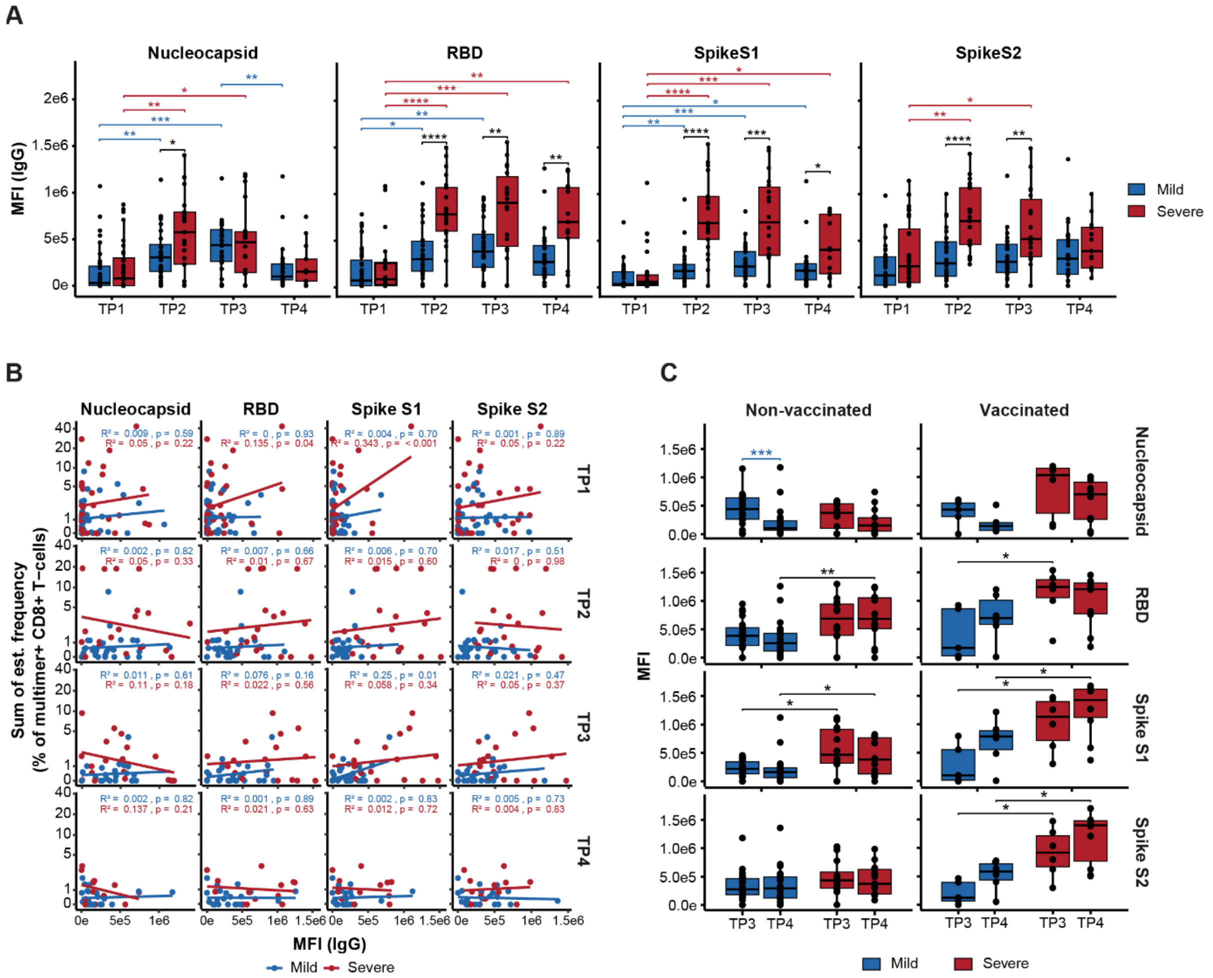

**Supplementary Fig. 4. SARS-CoV-2 antibody dynamics and correlation with T-cell responses in mild and severe COVID-19 patients.** (**A**) Levels of IgG antibodies against SARS-CoV-2 Spike protein subunits S1 and S2, the Spike receptor-binding domain (RBD), and nucleocapsid (N) protein in mild and severe COVID-19 patients from TP1 to TP4. (**B**) Correlation between the sum of the estimated frequencies (%) for the SARS-CoV-2 Spike-specific T-cell responses and the of IgG antibody levels in non-vaccinated COVID-19 patients across the four time points. The linear correlation coefficient ($r^2$) and p-values are indicated at the top of each plot. (**C**) Comparison between the levels of IgG antibodies against SARS-CoV-2 antigens for non-vaccinated and vaccinated mild and severe COVID-19 patients between TP3 and TP4. (A, C) Mann-Whitney test between disease severity and Mann-Whitney test adjusting p-values with the Bonferroni method for comparison between time points, **** ($p < 0.0001$), *** ($p < 0.001$), ** ($p < 0.01$) and * ($p \leq 0.05$).

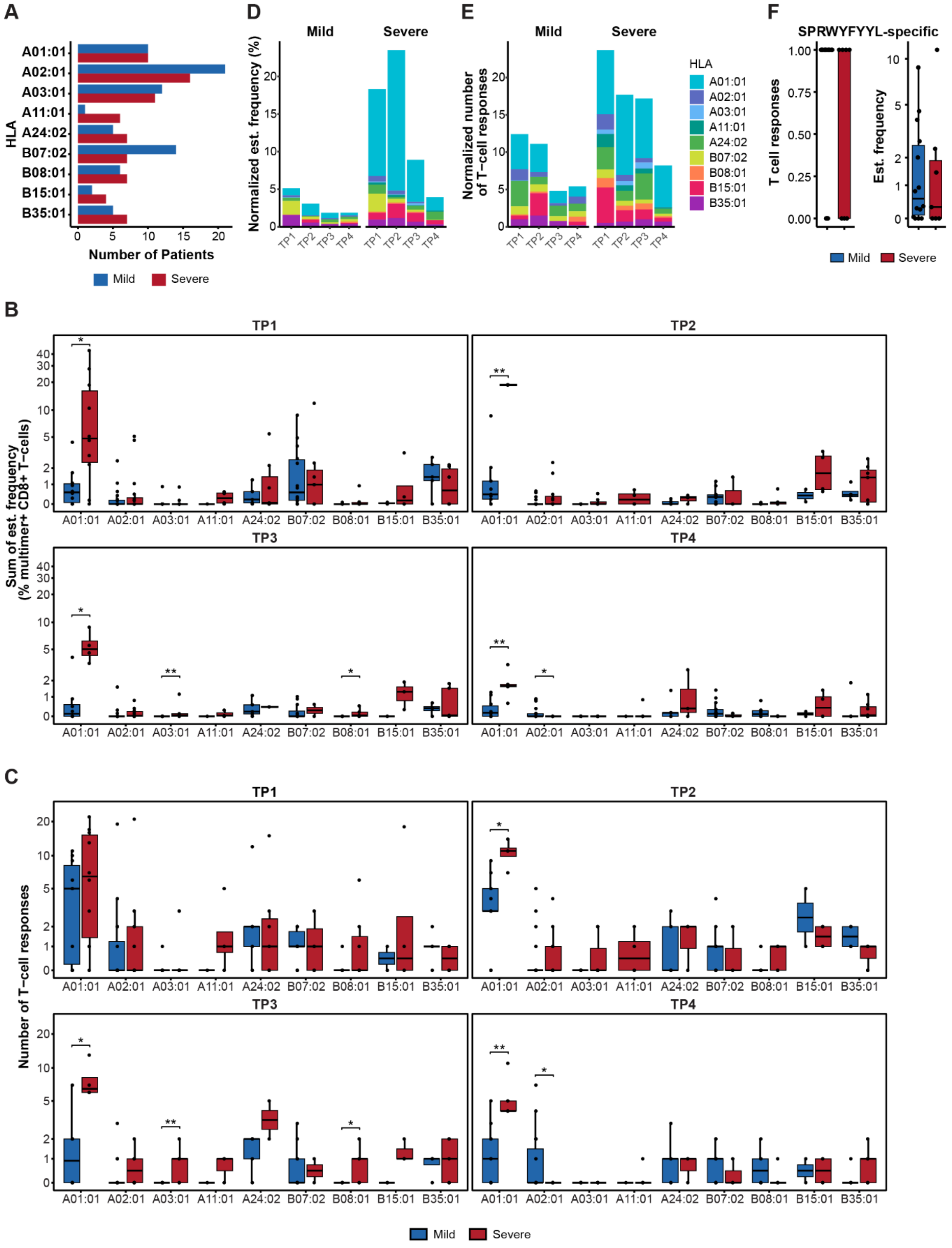


**Supplementary Fig. 5. HLA distribution of SARS-CoV-2-specific T-cell responses in mild and severe COVID-19 patients.** (**A**) Number of severe and mild patients specific to each HLA allele included in this study. Stacked bar plots summarize the normalized estimated frequency (**B**) and the normalized number (**C**) of SARS-CoV-2-specific T-cell responses per HLA allele, across four time points (TP1–TP4) in mild and severe patient cohorts. Normalization was performed based on the number of patients specific to each HLA allele at each time point for each disease severity. (**D**) Comparison of HLA-B07:02-restricted SPRWYFYYL-specific T-cell responses, represented by total responses and estimated frequencies (%), between mild and severe patients at TP1. Box plots comparing the sum of estimated frequencies (%) (**E**) and the total number (**F**) of SARS-CoV-2-specific T-cell responses restricted to each HLA allele between severe and mild COVID-19 patients across four time points post-diagnosis. (D-F) Mann-Whitney test, **** ($p < 0.0001$), *** ($p < 0.001$), ** ($p < 0.01$) and * ($p \leq 0.05$).

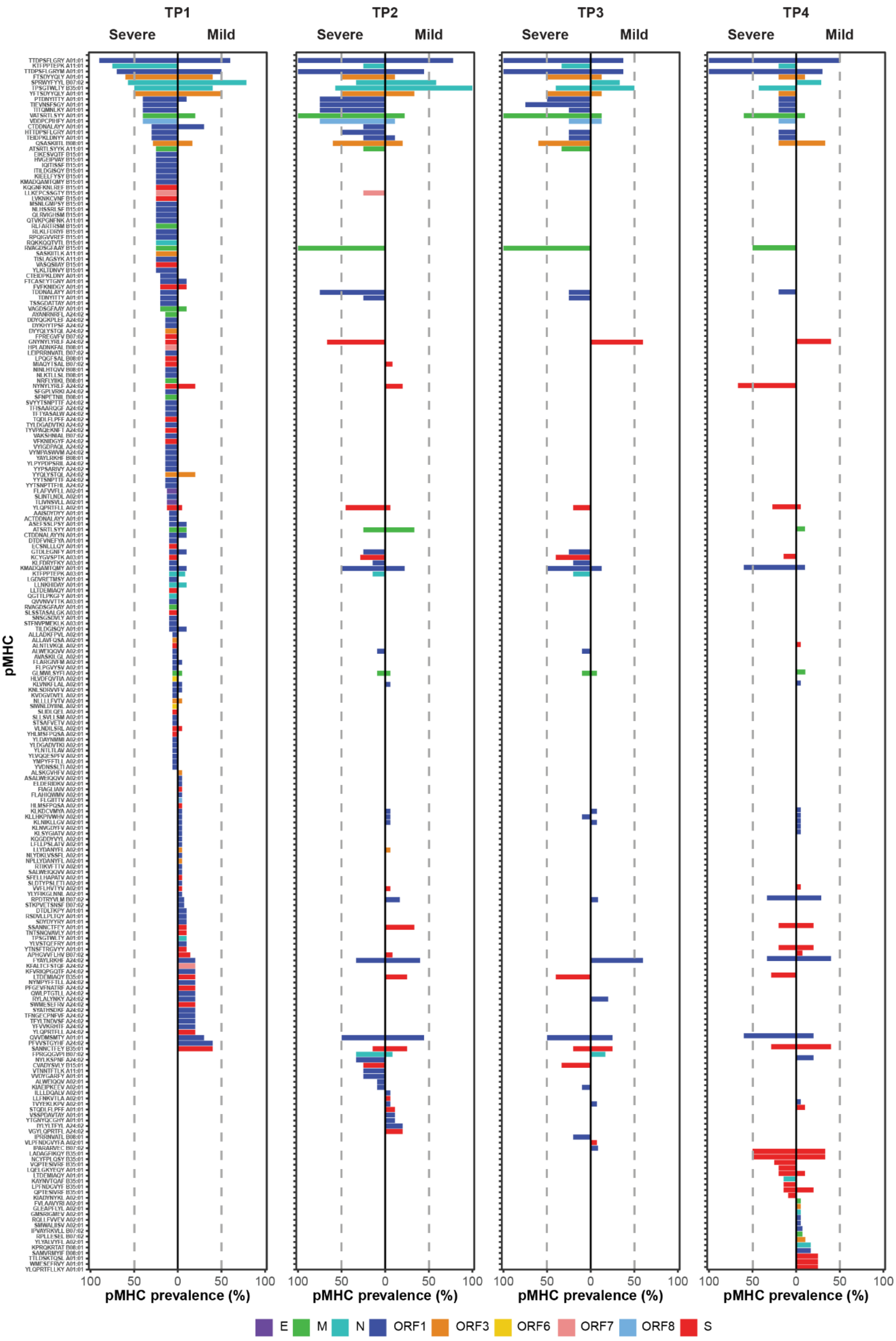


**Supplementary Fig. 6. Prevalence of CD8⁺ T-cell recognition towards SARS-CoV-2 epitopes.** Prevalence of T-cell recognition toward the individual epitopes detected in patients with COVID-19 split according to disease severity. TP4 prevalence includes T-cell responses towards additional Spike peptides included only in vaccinated patients. Only pMHC tested in more than 2 donors were included in this analysis. A dotted line is placed at 50% of prevalence to distinguish immunodominant epitopes. Bars are colored according to their protein of origin.

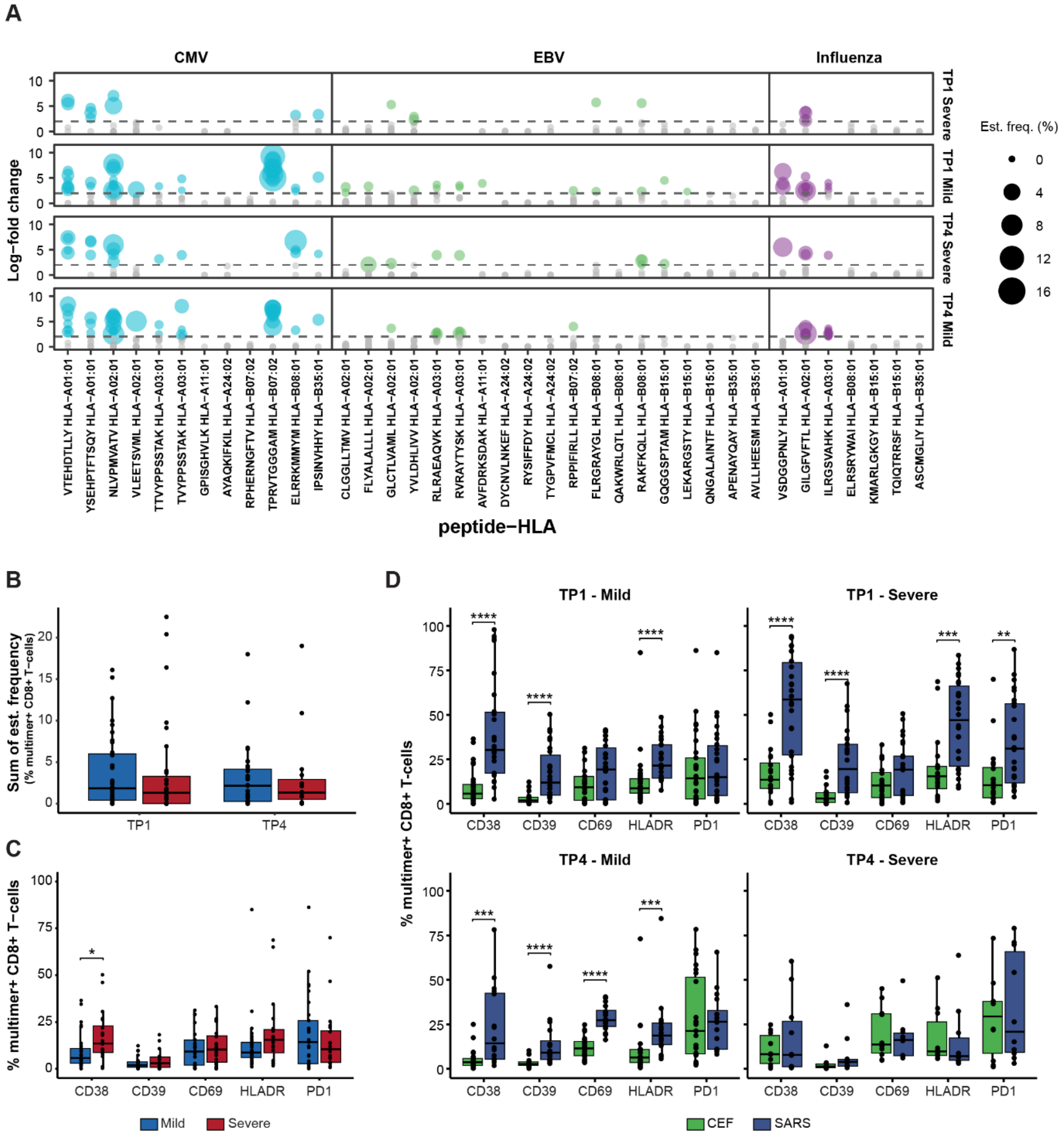


**Supplementary Fig. 7. Comparison of CEF- and SARS-CoV-2-specific T-cell responses in COVID-19 patients.** (**A**) CD8+ T-cell recognition to CEF-derived peptides at TP1 and TP4 in COVID-19 patients. Each dot represents one peptide-HLA combination per sample, and their size is proportional to their estimated frequency (%). (**B**) Box plot compares the sum of estimated frequency of CEF-specific T-cell epitopes between mild and severe COVID-19 patients at TP1 and TP4. Mann-Whitney test, no p-values were found to be significant. (**C**) Comparison of the percentage of CEF pMHC multimer+ CD8+ T-cells expressing the indicated surface markers between the mild and severe COVID-19 patients at TP1. (**D**) Box plot compares the percentage of SARS-CoV-2 pMHC multimer+ and CEF pMHC multimer+ CD8+ T-cells expressing the indicated surface markers in the COVID-19 patients at TP1 and TP4. Mann-Whitney test, **** ($p < 0.0001$), *** ($p < 0.001$), ** ($p < 0.01$) and * ($p \leq 0.05$).

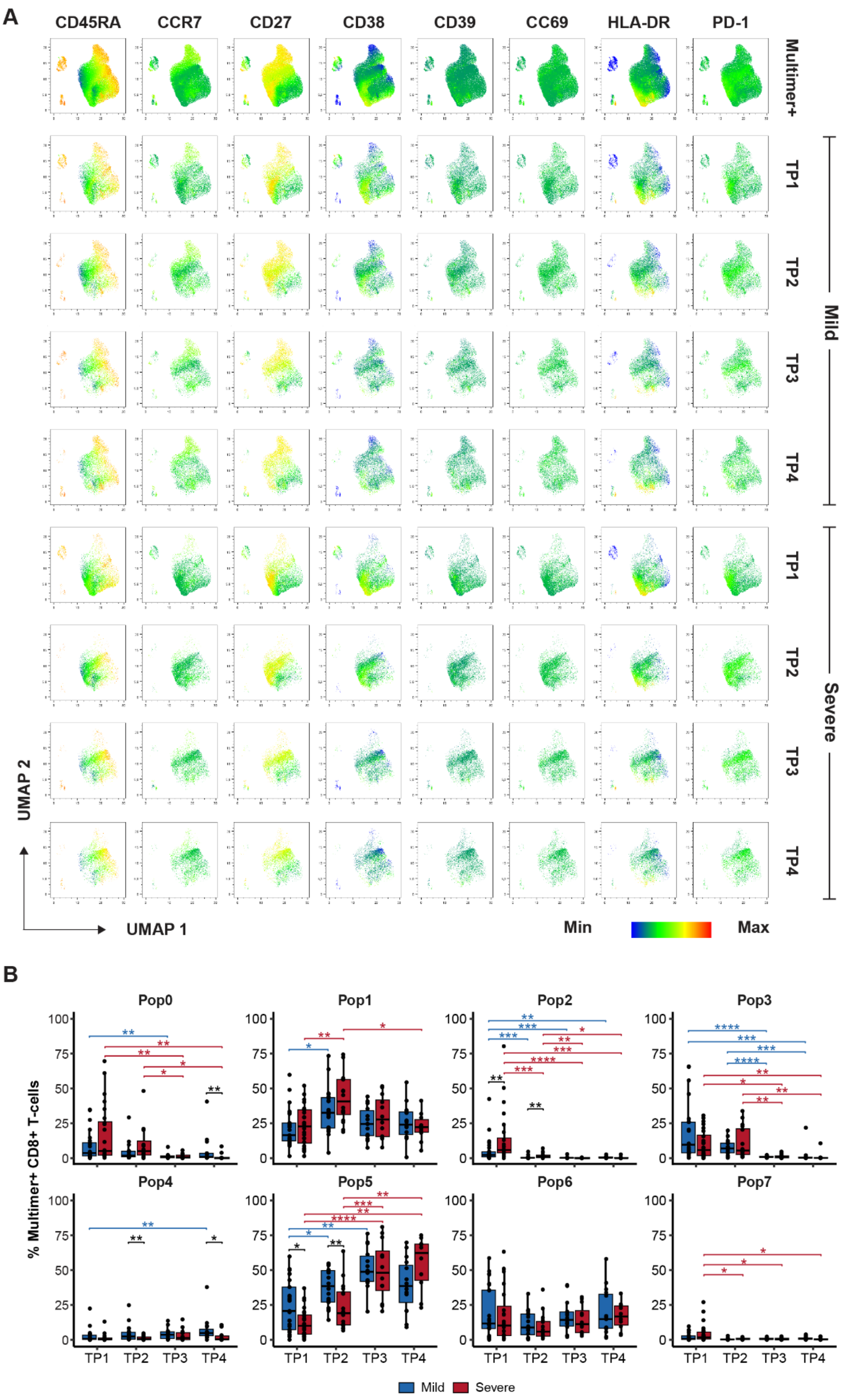


**Supplementary Fig. 8. SARS-CoV-2-specific CD8⁺ T-cells phenotype.** (**A**) UMAP showing the expression of individual phenotype markers CD45-RA, CCR7, CD27, CD38, CD39, CD69, HLA-DR, and PD-1 for SARS-CoV-2 pMHC multimer⁺ CD8⁺ T-cells in COVID-19 patients for all multimer⁺ T-cells (top row) and separated by severity and time point. (**B**) Box plot comparing the percentage of multimer positive CD8⁺ T-cells between mild and severe patients for each time point across all FlowSOM populations. Mann-Whitney test between disease severity and Mann-Whitney test adjusting p-values with the Bonferroni method for comparison between time points, **** ($p < 0.0001$), *** ($p < 0.001$), ** ($p < 0.01$) and * ($p \le 0.05$).

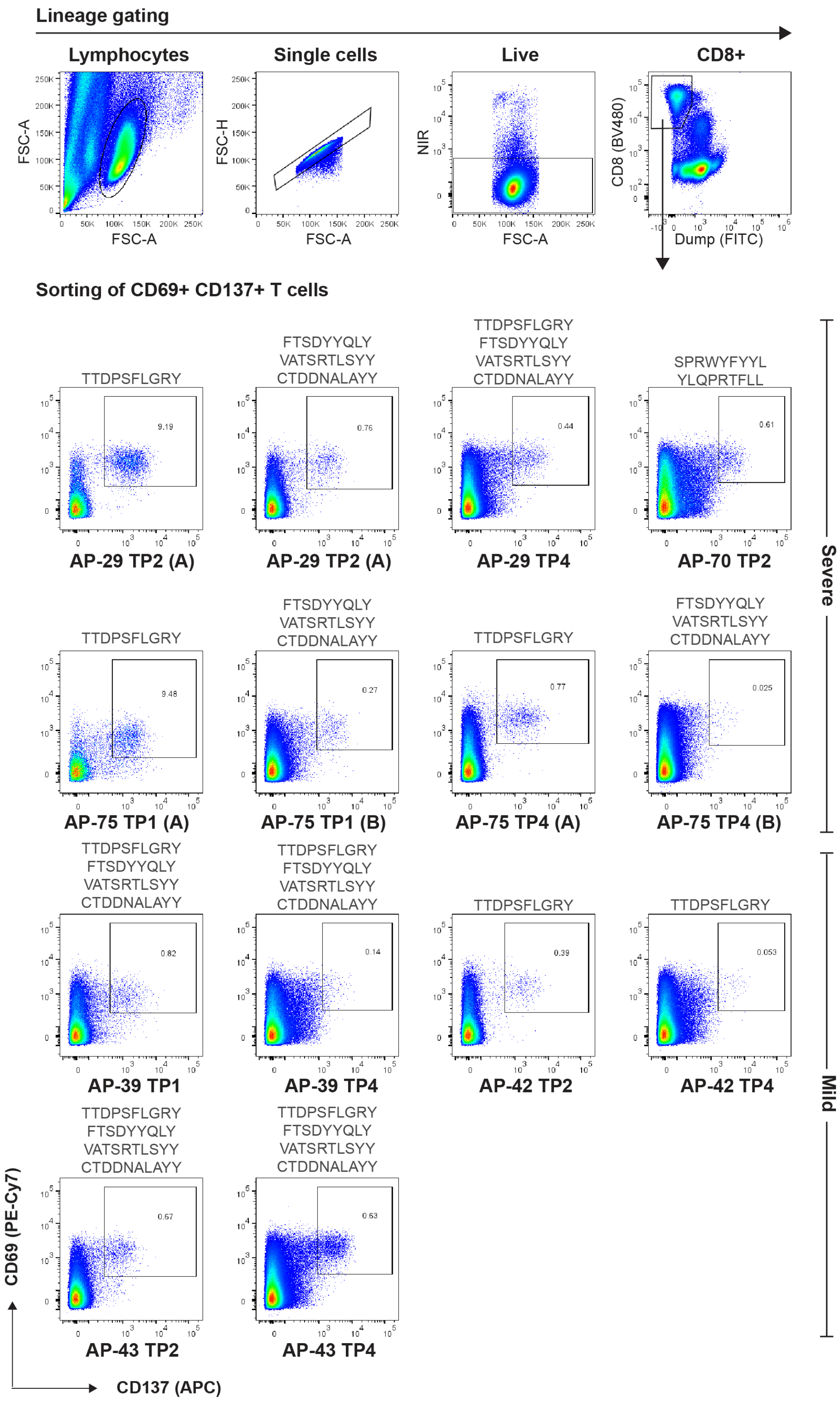


**Supplementary Fig. 9. Gating strategy for sorting of activated (CD69⁺ CD137⁺) CD8⁺ T-cells for single-cell analysis.** Representative flow cytometry plots showing the gating strategy on COVID-19 patient PBMCs to sort double positive CD69⁺ CD137⁺ T-cells used for single-cell analysis. Dot plots show the percentage of CD69⁺ CD137⁺ T-cell populations sorted from each sample.

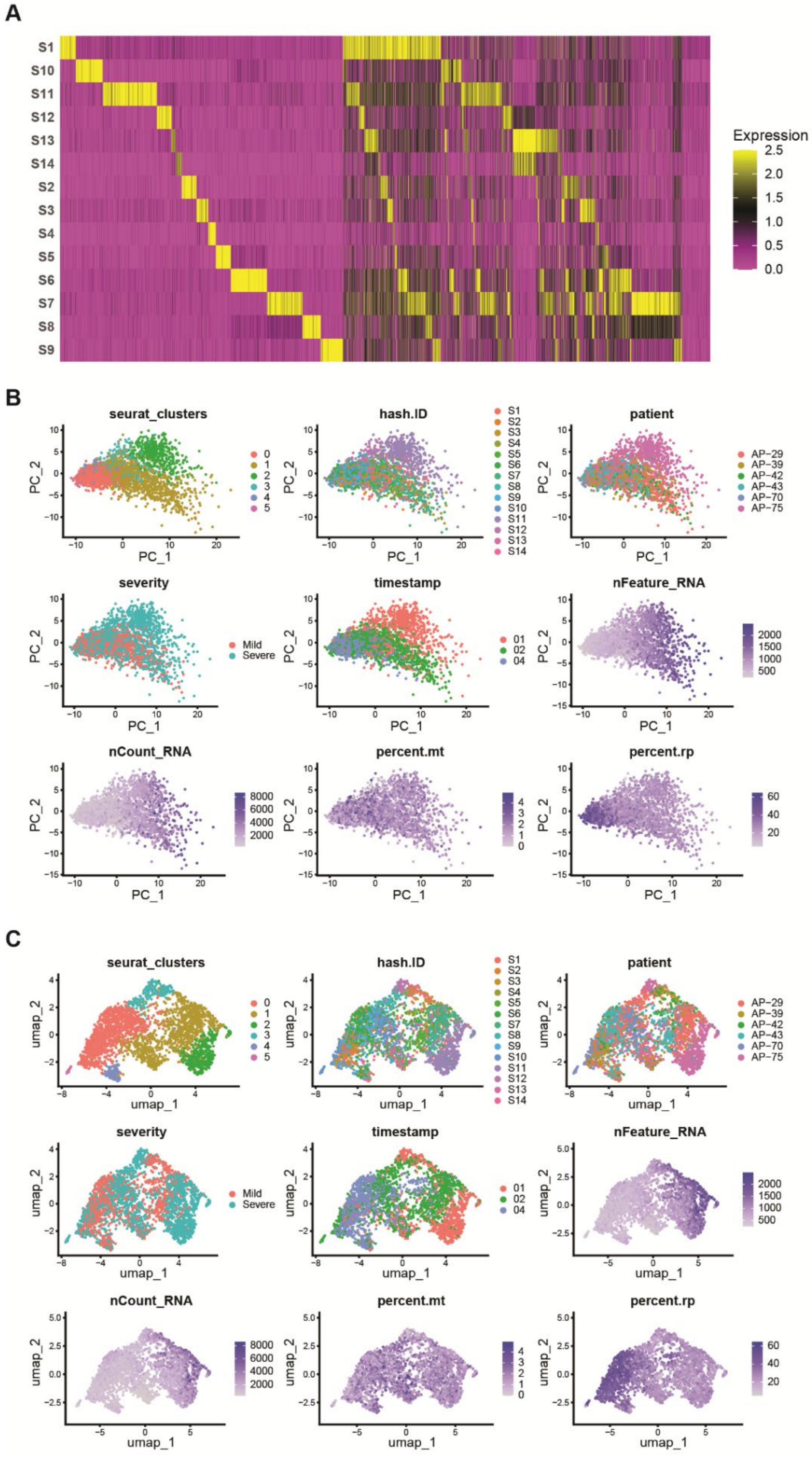


**Supplementary Fig. 10. Demultiplexing and single-cell transcriptome analysis.** (**A**) HTO heatmap after demultiplexing for all samples. (**B**) PCA plots for a selected set of variables. (**C**) UMAP plots for the selected set of variables. PCA: principal component analysis, HTO: hashtag oligo, UMAP: uniform manifold approximation and projection.

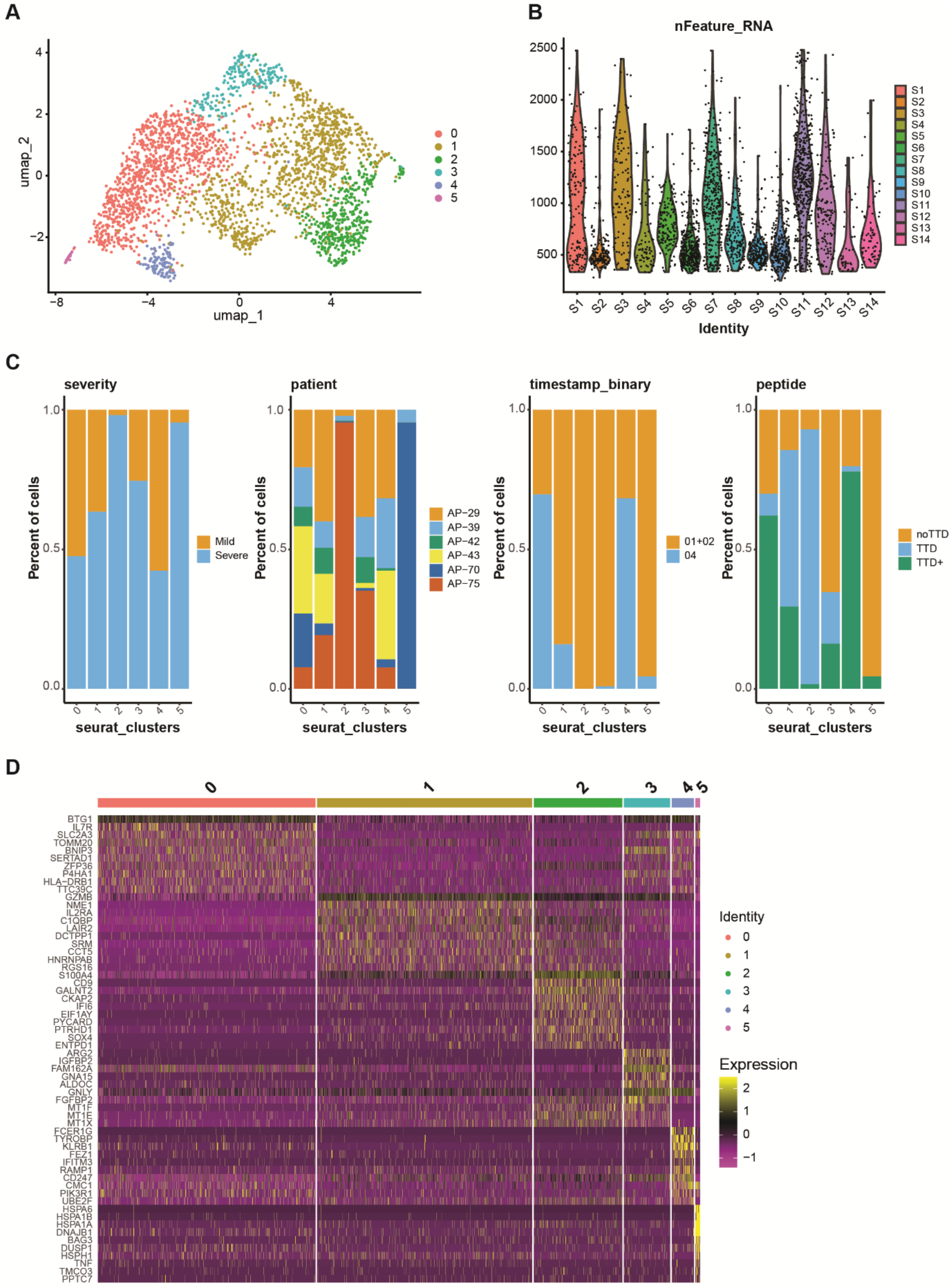


**Supplementary Fig. 11. Clinical and transcriptomic characteristics of single-cell clusters.** (**A**) UMAP of resulting clusters generated for all samples. (**B**) Distribution of nFeature_RNA across samples. (**C**) Distribution of clinical and experimental variables across clusters. (**D**) Heatmap of top 10 markers for each cluster.

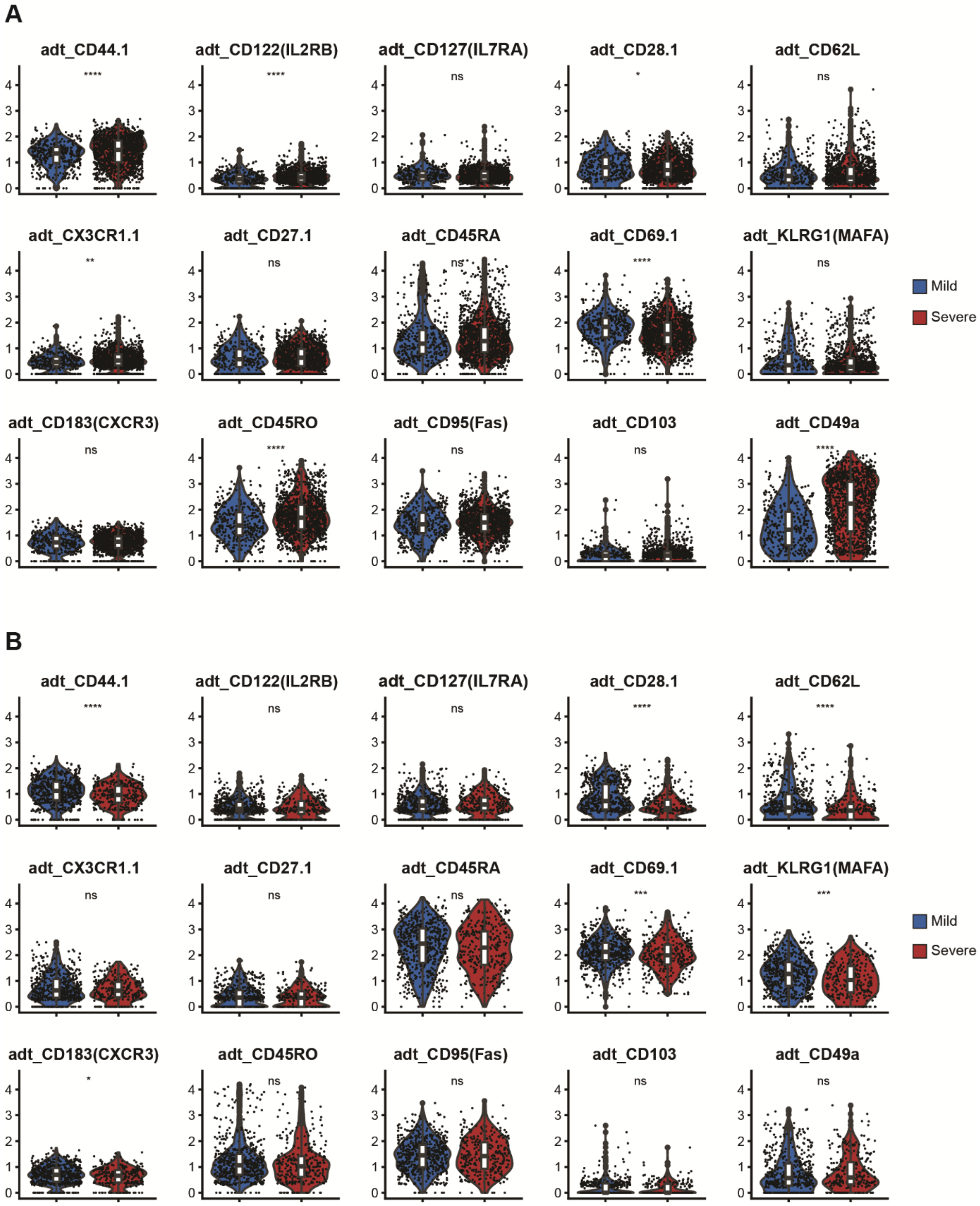


**Supplementary Fig. 12. Comparison of markers expression levels.** Violin plots comparing the expression levels of selected memory markers (surface markers) between mild and severe samples in early time point (**A**) and in late time point (**B**). Mann-Whitney test: *$p < 0.05$, **$p < 0.01$, ***$p < 0.001$, ****$p < 0.0001$.

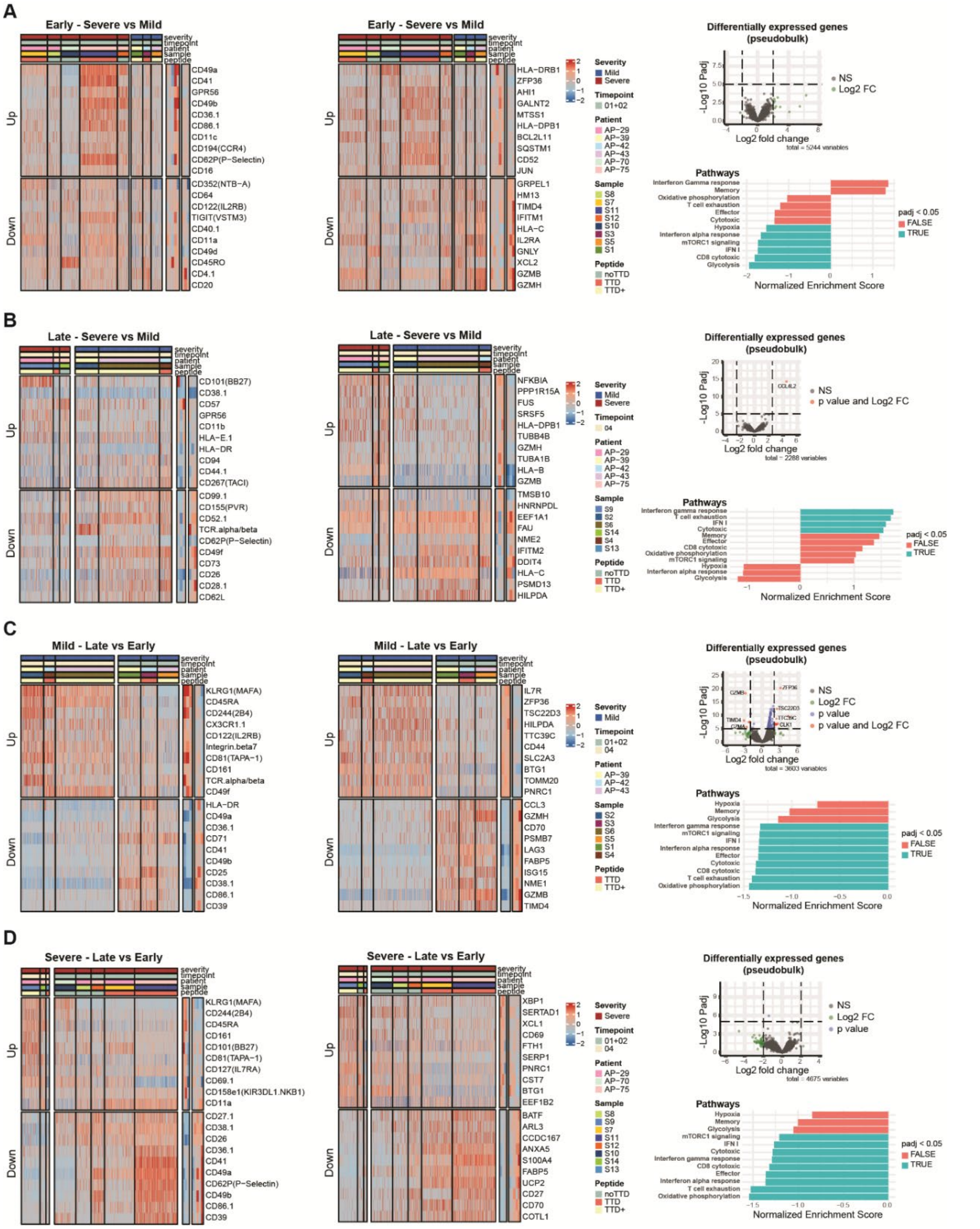


**Supplementary Fig. 13. Differential expression analysis of samples between different conditions.** (**A-D**) Differential expression analysis on a single-cell level for surface markers (**left heatmap**) and gene expressions (**right heatmap**). Top 10 markers and genes were selected from each side (avg_log2FC). Volcano plots (**right, top**) represent the results of pseudobulk differential expression gene analysis. Gene set enrichment analysis for selected gene sets (**right, bottom**). Comparison of severe vs mild patients in early (**A**) and late (**B**) COVID-19. Comparison of late vs early COVID-19 in mild (**C**) and severe (**D**) patients. TTD: TTDPSFLGRY; TTD+: TTDPSFLGRY + other peptides; noTTD: any peptide other than TTDPSFLGRY.

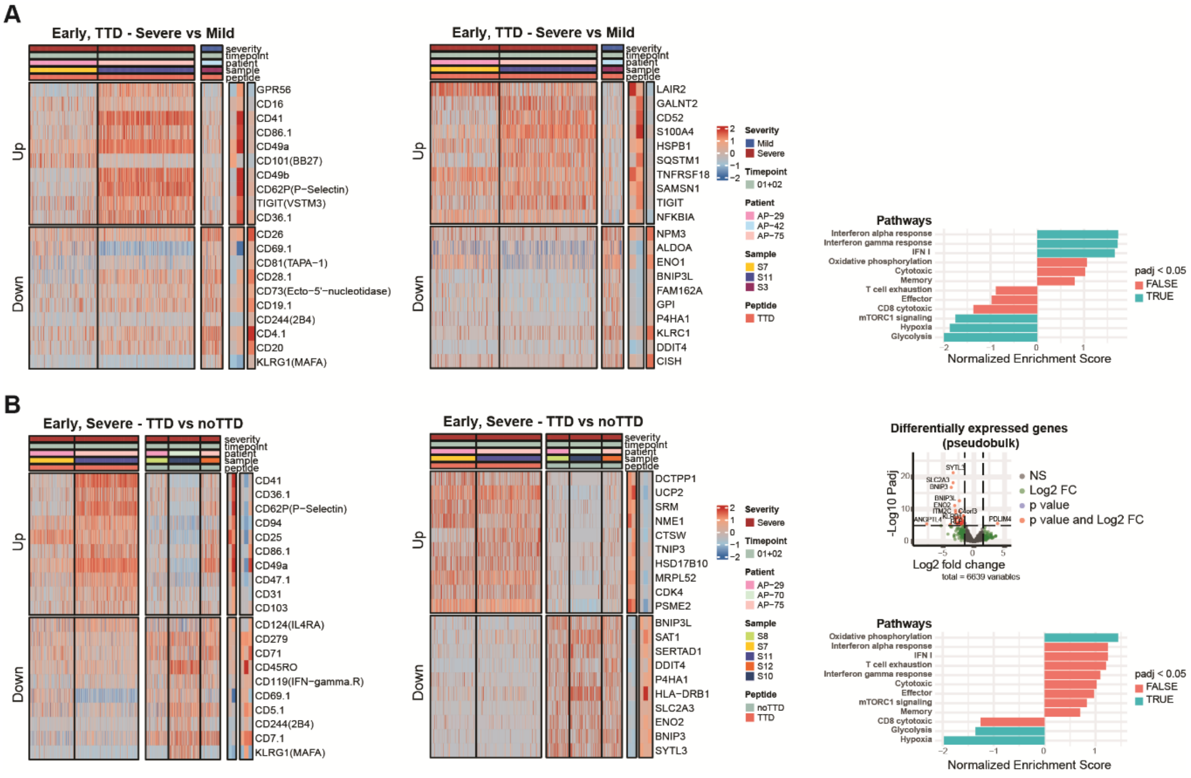


**Supplementary Fig. 14. Differential expression analysis in early (TP1/TP2) samples between different conditions.** (**A-B**) Differential expression analysis on a single-cell level for surface markers (**left heatmap**) and gene expressions (**right heatmap**). Top 10 markers and genes were selected from each side (avg_log2FC). Volcano plots represent results of pseudobulk differential expression gene analysis (**right, top**). Gene set enrichment analysis for selected gene sets (**right, bottom**). (**A**) Comparison of severe vs mild patients in early COVID-19, T-cells stimulated with TTD peptide. (**B**) Comparison of T-cells stimulated with TTD vs other peptides, in severe patients at late COVID-19. TTD: TTDPSFLGRY; noTTD: any peptide other than TTDPSFLGRY.

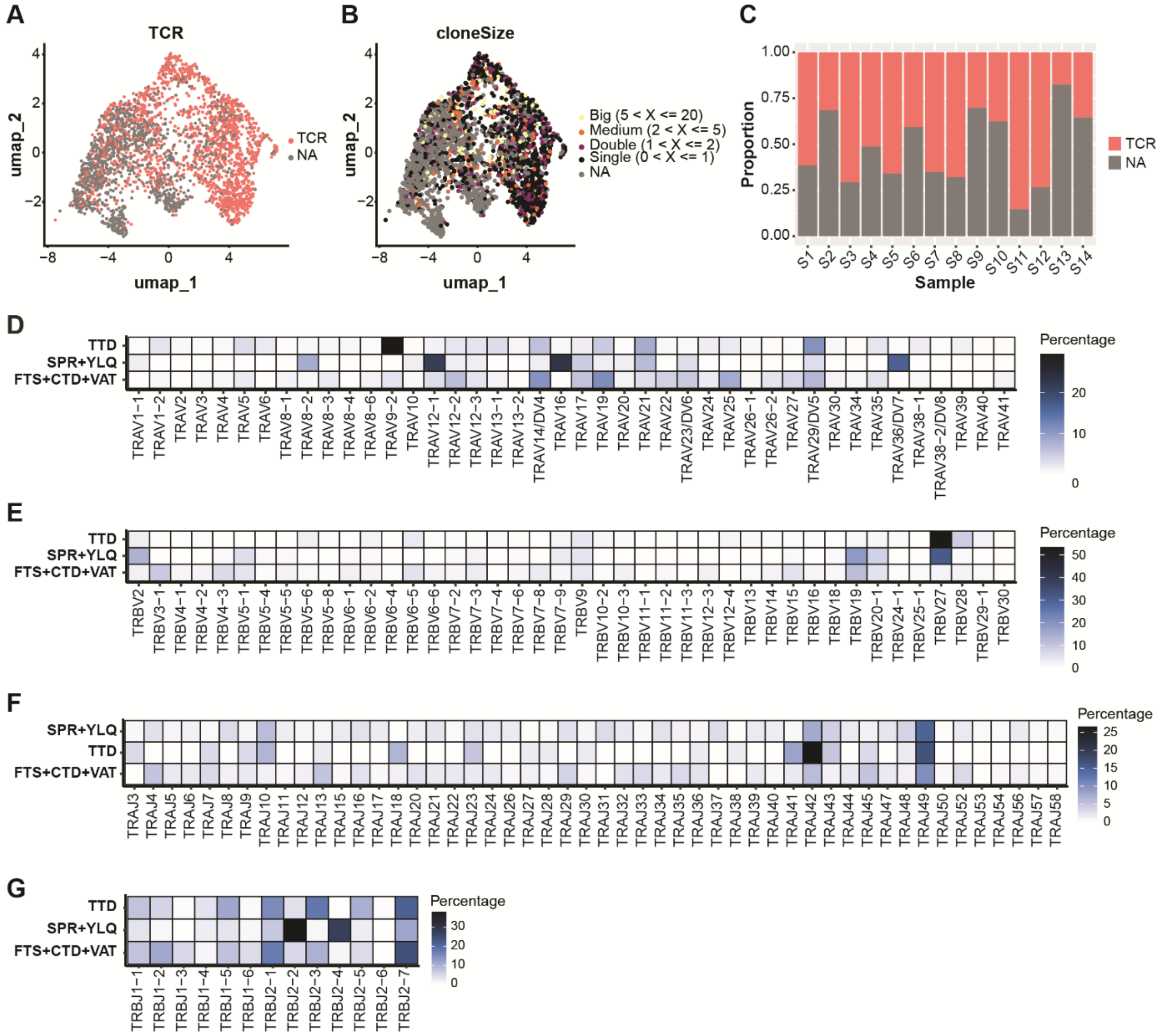


**Supplementary Fig. 15. Single-cell TCR sequencing analysis of SARS-CoV-2-specific responses. (A)** UMAP representation of all samples, colored by the availability of TCR information. (**B**) UMAP representation of all samples, colored by clone size. (**C**) Bar plot showing the distribution of TCR information availability across all samples. Heatmaps of the relative usage of V genes in alpha (**D**) and beta (**E**) chains for samples stimulated with various SARS-CoV-2 peptides. Heatmaps of the relative usage of J genes in alpha (**F**) and beta (**G**) chains for early time point samples stimulated with various SARS-CoV-2 peptides. TTD: TTDPSFLGRY, FTS: FTSDYYQLY, VAT: VATSRTLSYY, CTD: CTDDNALAYY, YLQ: YLQPRTFLL, SPR: SPRWYFYYL.

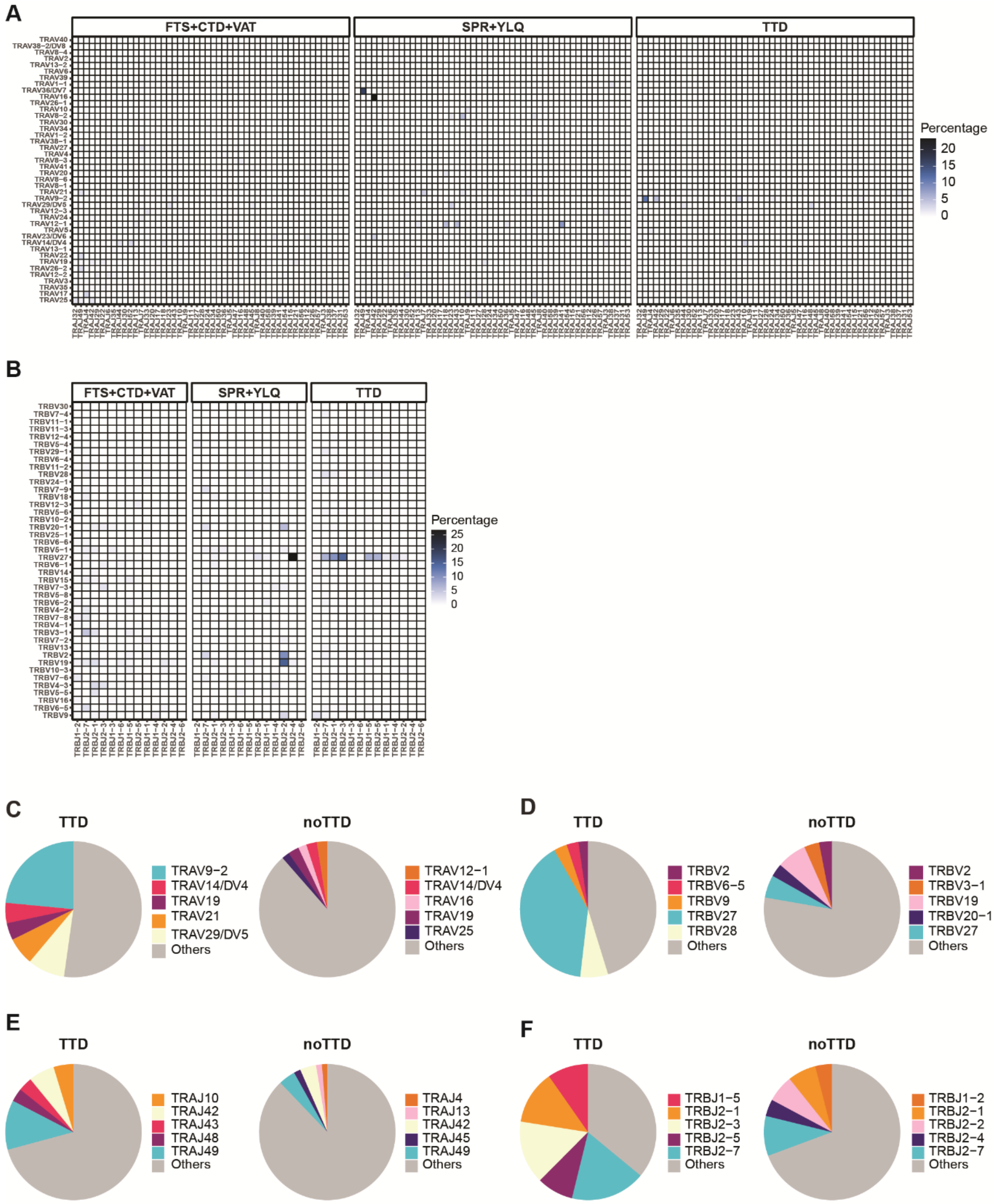


**Supplementary Fig. 16. Analysis of V and J gene usage in TCR alpha and beta chains for SARS-CoV-2-specific responses in early time points**. Heatmaps showing the relative usage of V and J gene combinations in the TCR alpha (**A**) and beta (**B**) chains for early-phase samples stimulated with SARS-CoV-2 peptides. Pie charts summarizing the top 5 most frequently used V genes in TTD versus no TTD samples for alpha (**C**) and beta (**D**) chains. Pie charts summarizing the top 5 J most frequently genes used in TTD versus no TTD samples at early time points for alpha (E) and beta (F) chains. TTD: TTDPSFLGRY, FTS: FTSDYYQLY, VAT: VATSRTLSYY, CTD: CTDDNALAYY, YLQ: YLQPRTFLL, SPR: SPRWYFYYL.

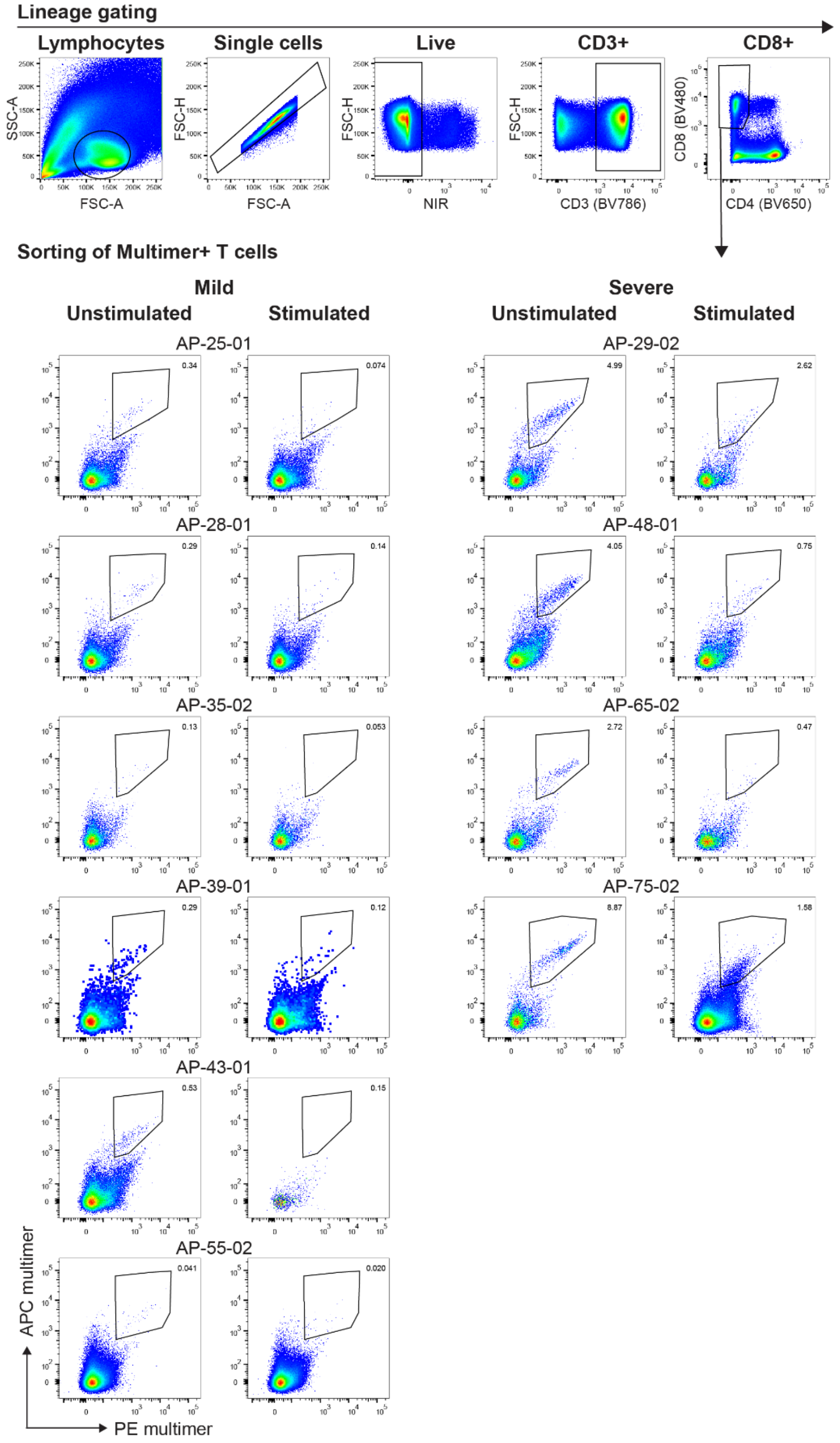


**Supplementary Fig. 17. Gating strategy for sorting of multimer⁺ CD8⁺ T-cells in Peptide-Stimulated COVID-19 Patients.** Representative flow cytometry plots demonstrating the gating strategy applied to PBMCs from COVID-19 patients (stimulated or unstimulated) for the identification and sorting of double-positive (PE⁺ APC⁺) multimer⁺ CD8⁺ T-cell populations. Dot plots display the percentages of PE⁺ APC⁺ multimer⁺ T-cells sorted from each patient sample (mild and severe COVID-19 cohorts). These sorted cells were used for analysis of TCR down-regulation following peptide stimulation.

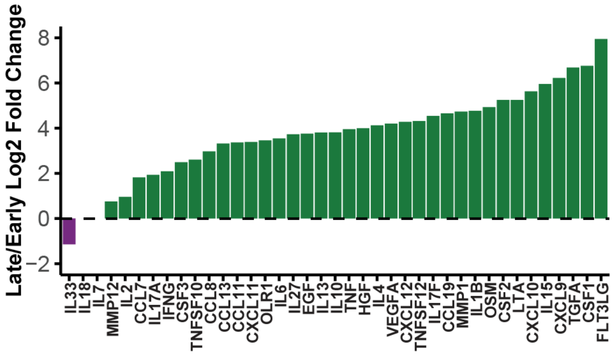


**Supplementary Fig. 18. Late-to-Early Protein Secretion Ratios in Severe COVID-19 Patients.** Bar plot showing the late-to-early ratio of the mean protein secretion levels, calculated using normalized protein concentrations across all peptides for each condition, in severe COVID-19 patients. Mann–Whitney test, no p-values were found to be significant.

### Supplementary Tables

**Supplementary Table 1.** Information on COVID-19 infected patients.

**Supplementary Table 2.** Timeline of blood sample collection.

**Supplementary Table 3.** SARS-CoV-2 epitopes included in longitudinal analysis with corresponding HLA rank scores.

**Supplementary Table 4.** SARS-CoV-2 peptides analyzed for T-cell reactivity with corresponding HLA rank scores.

**Supplementary Table 5.** Distribution of peptides used for longitudinal analysis across nine HLA-I allotypes and nine SARS-CoV-2 proteins.

**Supplementary Table 6.** HLA genotype data for COVID-19 infected donors.

**Supplementary Table 7.** Antibody panels.

**Supplementary Table 8.** CEF peptide library.

**Supplementary Table 9.** Summary of SARS-CoV-2-specific T-cell responses in COVID-19 patients.

**Supplementary Table 10.** SARS-CoV-2-specific T-cell epitopes identified in COVID-19 patients across four time points, including Spike responses in vaccinated individuals between TP3 and TP4.

**Supplementary Table 11.** Statistical comparison of SARS-CoV-2-specific T-cell responses between mild and severe COVID-19 patients.

**Supplementary Table 12.** Prevalence of SARS-CoV-2-specific peptides tested in at least three individuals sharing the corresponding HLA molecule, by time point and severity group.

**Supplementary Table 13.** Statistical significance of differences in the number of T-cell responses between mild and severe patients.

**Supplementary Table 14.** Additional SARS-CoV-2 Spike-derived peptides used for analysis of T-cell reactivity at TP4 in COVID-19 vaccinated patients.

**Supplementary Table 15.** CEF-specific T-cell epitopes identified in COVID-19 patients at TP1 and TP4.

**Supplementary Table 16.** Experimental details for single-cell analysis.

**Supplementary Table 17.** SARS-CoV-2 peptides used for TCR downregulation analysis.

**Supplementary Table 18.** SARS-CoV-2 peptides used for single-peptide PBMC stimulation for Olink analysis.
